# CBP/P300 BRD Inhibition Reduces Neutrophil Accumulation and Activates Antitumor Immunity in TNBC

**DOI:** 10.1101/2024.04.25.590983

**Authors:** Xueying Yuan, Xiaoxin Hao, Hilda L. Chan, Na Zhao, Diego A. Pedroza, Fengshuo Liu, Kang Le, Alex J. Smith, Sebastian J. Calderon, Nadia Lieu, Michael J. Soth, Philip Jones, Xiang H.-F. Zhang, Jeffrey M. Rosen

## Abstract

Tumor-associated neutrophils (TANs) have been shown to promote immunosuppression and tumor progression, and a high TAN frequency predicts poor prognosis in triple-negative breast cancer (TNBC). Dysregulation of CREB binding protein (CBP)/P300 function has been observed with multiple cancer types. The bromodomain (BRD) of CBP/P300 has been shown to regulate its activity. In this study, we found that IACS-70654, a novel and selective CBP/P300 BRD inhibitor, reduced TANs and inhibited the growth of neutrophil-enriched TNBC models. In the bone marrow, CBP/P300 BRD inhibition reduced the tumor-driven abnormal differentiation and proliferation of neutrophil progenitors. Inhibition of CBP/P300 BRD also stimulated the immune response by inducing an IFN response and MHCI expression in tumor cells and increasing tumor-infiltrated CTLs. Moreover, IACS-70654 improved the response of a neutrophil-enriched TNBC model to docetaxel and immune checkpoint blockade. This provides a rationale for combining a CBP/P300 BRD inhibitor with standard-of-care therapies in future clinical trials for neutrophil-enriched TNBC.

**Summary:** In neutrophil-enriched triple-negative breast cancer (TNBC) models, CREB binding protein (CBP)/P300 bromodomain (BRD) inhibition reduces tumor growth and systemic neutrophil accumulation while stimulating an antitumor immune response. This improves standard-of-care therapies, suggesting a potential therapeutic benefit of CBP/P300 BRD inhibitors for neutrophil-enriched TNBC.

## Introduction

Triple-negative breast cancer (TNBC) is a biologically heterogeneous and clinically important breast cancer subtype defined by the lack of estrogen receptor, progesterone receptor, and human epidermal growth factor receptor 2 amplification (Bianchini et al., 2016). In the tumor immune microenvironment (TIME) of TNBC, tumor-associated myeloid cells including tumor-associated neutrophils (TANs) and tumor-associated macrophages (TAMs) are the most abundant infiltrated immune cells (Gentles et al., 2015, Wu and Zhang, 2020). TANs contribute to tumor progression through T cell inhibition, promoting tumor proliferation and therapy resistance (Wu and Zhang, 2020, Keeley et al., 2019). TNBC exhibits heterogeneous frequencies of TANs, and high infiltration of TANs has been associated with poor prognosis (Kim et al., 2019, Gentles et al., 2015). Moreover, systemic changes such as the accumulation of blood neutrophils and overproduction of immature myeloid cells in the bone marrow have been observed in TNBC (Kim et al., 2019, Casbon et al., 2015, Hao et al., 2023, Veglia et al., 2018). TANs can also facilitate tumor metastasis by inducing invasion, migration, and epithelial-mesenchymal transition (EMT) (Keeley et al., 2019, Wu and Zhang, 2020).

Epigenetic modifications have been demonstrated to reprogram the functions and accumulation of TANs (Xu et al., 2022, Lodewijk et al., 2021). Previously, we performed an organoid screen in search of epigenetic inhibitors that can reverse EMT, and IACS-70654, a novel and selective CREB binding protein (CBP)/P300 bromodomain (BRD) inhibitor, was identified as one of the top hits in this screen (Zhao et al., 2021). CBP and P300 are transcriptional coactivators that are highly homologous, especially in sequences encoding histone acetyltransferase (HAT) domains and BRDs (Zeng et al., 2008, Dancy and Cole, 2015). The HAT domain of CBP/P300 can acetylate histones and some transcription factors (e.g., p53) to regulate gene expression (Dancy and Cole, 2015, Wang et al., 2013). Functional dysregulation of CBP/P300 has been associated with tumorigenesis (Wang et al., 2013). It has been difficult to develop highly selective inhibitors directly targeting their HAT domains, and thus these have not progressed to the clinic. However, inhibitors targeting the BRDs have shown promising results (Breen and Mapp, 2018). The BRD of CBP/P300 is involved in the positive feedback of histone acetylation and transcription activation (Das et al., 2014, Kikuchi et al., 2023). CBP/P300 BRD inhibitor CCS147 is in clinical trials for hematologic malignancies (Nicosia et al., 2023, Raisner et al., 2018). More importantly, inhibition of P300/CBP BRD was able to reduce the growth of a human TNBC xenograft model MDA-MB-231 X1.1 and reprogrammed the tumor-associated myeloid cells (de Almeida Nagata et al., 2019). However, the effects of CBP/P300 BRD inhibition on TNBC and the TIME have not been thoroughly investigated in immunocompetent models.

Therefore, this study was designed to test the effects of the novel CBP/P300 BRD inhibitor IACS-70654 on both tumor cells and the TIME of multiple immunocompetent mouse models of TNBC with heterogeneous TAN and TAM frequencies. We found that IACS-70654 inhibited the growth of neutrophil-enriched TNBC models and reduced immunosuppression by decreasing TANs and inhibiting abnormal neutrophil production in the bone marrow. IACS-70654 also exerted immune stimulation by inducing both an interferon (IFN) and a T-cell response.

## Results

### IACS-70654 inhibited the growth of neutrophil-enriched syngeneic mammary tumor models

To examine the effects of CBP/P300 BRD inhibition on TNBC *in vivo*, we selected four immunocompetent mouse models derived from either BALB/c or C57BL/6 background. 2208L, T6, and T12 tumors are syngeneic *Trp53*-null TNBC models, which were shown to recapitulate the aggressiveness, heterogeneity, and resistance to standard-of-care therapies in human TNBC (Herschkowitz et al., 2012, Gerber-Ferder et al., 2023, Grieshaber-Bouyer et al., 2021). PyMT-N is a luminal-like subtype derived from MMTV-PyMT tumors, exhibiting stable myeloid cell infiltration and resistance to immune checkpoint blockade (ICB) (Kim et al., 2019). These models displayed distinct TAN and TAM frequencies. TANs are defined as CD45^+^CD11b^+^Ly6G^+^Ly6C^med-low^ in flow cytometry and S100A8^+^ in immunostaining. TAMs are defined as CD45^+^CD11b^+^Ly6G^-^Ly6C^-^F4/80^+^ in flow cytometry and F4/80^+^ in immunostaining.

Based on flow cytometry, immunostaining, and previous publications, 2208L, PyMT-N, and T6 can be categorized as neutrophil-enriched models, and T12 can be categorized as a macrophage-enriched model (Singh et al., 2022, Kim et al., 2019) (**Figure 1A and B**). Within the neutrophil-enriched models, PyMT-N and T6 tumors were infiltrated with more TAMs than 2208L tumors (**Figure 1B**). Tumor pieces (*Trp53-null* models) or freshly dissociated tumor cells (PyMT-N) were implanted into the mammary fat pad of BALB/c or C57BL/6 mice. The mice were randomized and treated with vehicle or IACS-70654 when the tumors were palpable and reached more than 80 mm^3^ in volume on average (**Figure 1C**). IACS-70654 is a novel, potent, and highly selective inhibitor of CBP/P300 BRD (Supplemental Table S1 and 2 and Figure S1A). IACS-70654 was orally administered at the dosage of 3.75 mg/kg on a 3-on/2-off regimen for all animal studies, and therefore a total of 6 doses were given in a 7-day experiment (**Figure 1C**).

**Figure 1.**
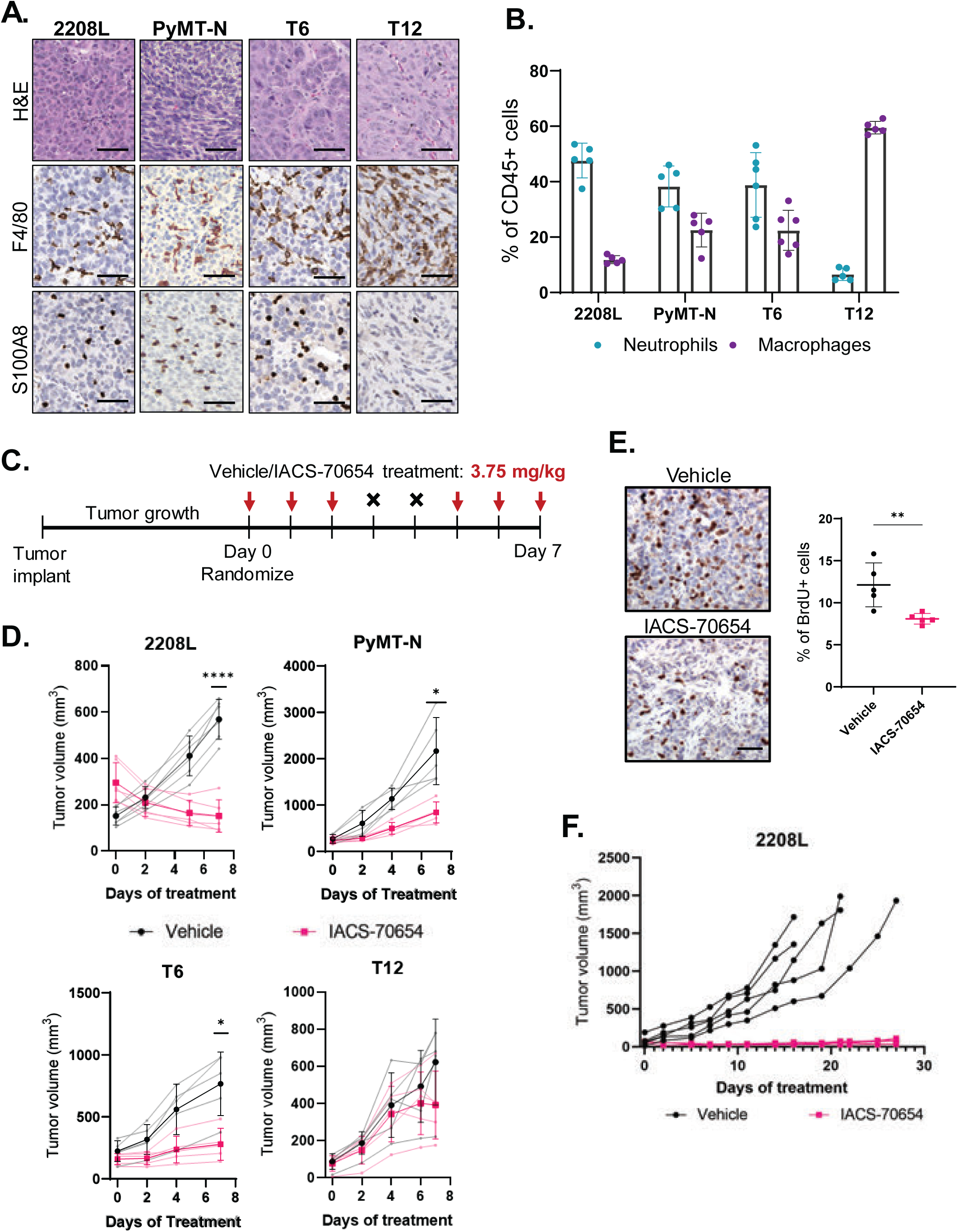
IACS-70654 suppressed the growth of neutrophil-enriched preclinical mouse models of TNBC. **A.** Representative images of H&E and immunostaining of all TNBC preclinical mouse models used. F4/80 is a macrophage marker, and S100A8 is a neutrophil marker. Scale bars: 50 µm. **B.** Flow cytometry analysis of tumor-associated neutrophils and macrophages in all TNBC preclinical mouse models used (*n* ≥ 5 for all models). Error bars represent SD. **C.** Treatment schemes of IACS-70654 in a 7-day experiment. A small piece from a fresh *Trp53*-null tumor (2208L, T6, or T12) was implanted into the fourth left mammary fat of each BALB/c mouse. For the PyMT-N model, 250,000 freshly dissociated cells were injected into the same position of each C57BL/6 mouse. When the average tumor volume reached 80-250 mm^3^, the mice were randomized, and then the treatment was initiated. Vehicle or IACS-70654 at 3.75 mg/kg was administered orally on a 3-on/2-off schedule. On Day 7, mice were euthanized 2 hrs after receiving the last treatment. **D.** Tumor growth curves of the *Trp53*-null (2208L, T6, and T12) and PyMT-N tumors treated with IACS-70654 for 7 days. For 2208L, *n* = 6, and for all other models, *n* = 5. Two-way ANOVA and Sidak’s multiple comparisons test were used. *, p<0.05; ****, p<0.0001. Error bars represent SD. **E.** Left: Representative images of immunostaining of BrdU in 2208L tumor sections treated with vehicle or IACS-70654 for 7 days. Scale bar: 50 µm. Right: Quantification of BrdU staining using Fiji. Three representative images were analyzed for each tumor and *n* = 5 biological replicates for each treatment arm. Two-tailed unpaired Student’s *t* test was used. **, p<0.01. Error bars represent SD. **F.** Tumor growth curves of 2208L tumors treated with IACS-70654 for 27 days.

IACS-70654 treatment was well-tolerated and did not lead to significant body weight loss (Supplemental Figure S1B). Surprisingly IACS-70654 as a single agent resulted in the regression of 2208L tumors, which has the highest TAN frequency (**Figure 1B and D** and Supplemental Figure S1C). IACS-70654 reduced the growth of PyMT-N and T6 tumors, which are neutrophil-enriched but infiltrated with more TAMs than 2208L tumors (**Figure 1B and D** and Supplemental Figure S1C). The macrophage-enriched T12 model was resistant (**Figure 1B and D**, Supplemental Figure S1C). Tumor regression is rarely observed with a single-agent treatment in *Trp53-null* models because they are highly aggressive and resistant to therapies.

We also used BrdU to determine the effect of IACS-70654 on tumor cell proliferation. 2208L tumors treated with IACS-70654 showed a reduction in BrdU incorporation and thus were less proliferative (**Figure 1E**). To test whether tumor inhibition can persist in long-term treatment, we conducted a 27-day treatment study of IACS-70654 with 2208L tumors. IACS-70654 durably inhibited the growth of 2208L tumors, and tumors showed no signs of resistance (**Figure 1F**).

Based on those findings, we hypothesized that the infiltrated myeloid cells might explain the difference in response. Therefore, we next examined the changes in infiltrated myeloid cells after IACS-70654 treatment.

### IACS-70654 reduced TANs and immunosuppression in the TIME

To investigate the effects of IACS-70654 on infiltrated myeloid cells, we analyzed the infiltrated immune cells of the selected models from the 7-day treatment study. IACS-70654 reduced Ly6G expression in the TANs of all 4 models and significantly reduced the percentage of TANs in 2208L, PyMT-N, and T12 tumors (**Figure 2A** and Supplemental Figure S2A). The reduction in TANs was verified with reduced immunostaining of S100A8 in 2208L tumors and was observed within 72 hrs after starting treatment (**Figure 2B** and Supplemental Figure S2B). The TANs of neutrophil-enriched models were demonstrated to be immunosuppressive and categorized as granulocytic myeloid-derived suppressor cells (gMDSCs), but TANs of macrophage-enriched tumors resembled normal neutrophils (Kim et al., 2019). Therefore, reduced TANs should not reduce immunosuppression in T12 tumors, and thus this may partially explain why T12 did not respond to IACS-70654. Higher TAM frequency in PyMT-N and T6 might explain why regression was only observed in 2208L tumors. TANs in 2208L and PyMT-N tumors treated with IACS-70654 were also found to have reduced expression of histone H3 lysine 27 acetylation (H3K27ac), the target biomarker for IACS-70654 (**Figure 2C**). This result suggested that TANs can be directly reprogrammed by IACS-70654. Moreover, high TAN frequency in tumors is accompanied by an accumulation of peripheral blood neutrophils (Kim et al., 2019, Casbon et al., 2015). IACS-70654 reduced blood neutrophils in 2208L tumor-bearing mice to a similar level as non-tumor-bearing mice (**Figure 2D**). We also treated non-tumor-bearing mice with IACS-70654 and observed no significant change in blood neutrophil (Supplemental Figure S2C).

**Figure 2.**
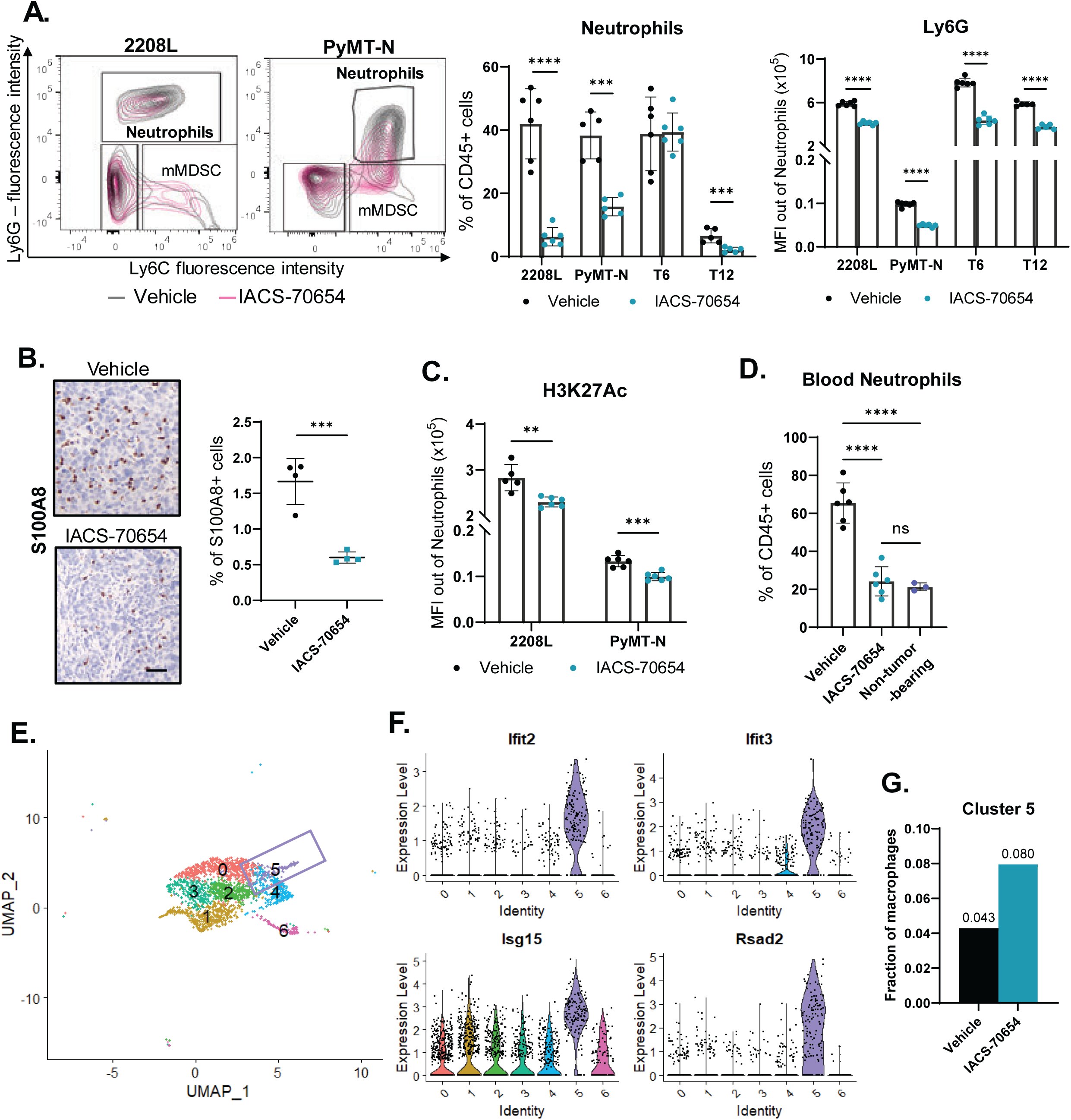
IACS-70654 reduced TANs and reprogrammed myeloid cell compositions in the TIME. **A.** Flow cytometry analyses of the infiltrated immune cells in 2208L, PyMT-N, T6, and T12 tumors treated with vehicle or IACS-70654 for 7 days. Left: Representative contour plots of Ly6G versus Ly6C showing gating strategy for and changes in myeloid populations. The gating was performed on CD45^+^/CD11b^+^ populations. Middle: Quantification of TANs as percentages of CD45+ cells. Neutrophils are defined by Ly6G^+^/Ly6C^med-low^. Right: Medium fluorescent intensity (MFI) of Ly6G in the TANs. Two-tailed unpaired Student’s *t* test was used. ***, p<0.001; ****, p<0.0001. For all models, at least five biological replicates were used for each treatment arm. Error bars represent SD. **B.** Left: Representative images of immunostaining of S100A8 on sections of 2208L tumor treated with vehicle or IACS-70654 for 7 days. Scale bar: 50 µm. Right: Quantification of S100A8 staining using Fiji. Three representative images were analyzed for each tumor, and four biological replicates were used for each treatment arm. Two-tailed unpaired Student’s *t* test was used. ***, p<0.001. Error bars represent SD. **C.** Median fluorescent intensity (MFI) of H3K27ac in flow cytometry analyses of TANs in 2208L and PyMT-N tumors treated with vehicle or IACS-70654. For all models, at least five biological replicates were used for each treatment arm. Error bars represent SD. **D.** Flow cytometry analyses of blood neutrophils in 2208L tumor-bearing mice treated with vehicle or IACS-70654 and untreated non-tumor-bearing BALB/c mice. Ordinary one-way ANOVA and Tukey’s multiple comparisons test were used. Five biological replicates were used for each treatment arm of 2208L tumor-bearing mice, and three biological replicates were used for non-tumor-bearing mice. ****, p<0.0001; ns, p>0.05. Error bars represent SD. **E.** UMAP plot TAM population (monocytes included) with clustering performed at the resolution of 0.5. **F.** Violin plot showing the expression of representative IFN-associated genes (*Ifit2, Ifit3, Isg15, Rsad2*) across all clusters of the TAM population. **G.** Fraction of Cluster 5 in the TAM population. The fraction value is labeled.

Besides neutrophils, IACS-70654 also has effects on other infiltrated myeloid cells. Even in neutrophil-enriched tumors, TAMs remain the second most abundant immune cell, indicating their importance (Kim et al., 2019) (**Figure 1B**). TAMs can have various functions, leading to an antitumor or protumor phenotype (Ma et al., 2022). To determine transcriptional changes in TAMs, single-cell RNA sequencing (scRNA-seq) was performed on 2208L tumors treated with vehicle and IACS-70654 for 7 days, and the TAM cluster was isolated. Six clusters with distinct RNA expression profiles were identified in TAMs of 2208L tumors (**Figure 2E** and Supplemental Figure S2D and E). Cluster 5 of TAMs highly expressed IFN-response genes (e.g. *Ifit2, Ifit3, Isg15, Rsad2*), which are associated with the antitumor phenotype (Mehta et al., 2021). IACS-70654 treatment led to an almost 2-fold increase in the fraction of cluster 5. Moreover, monocytes are defined as CD45^+^CD11b^+^Ly6G^-^Ly6C^+^, and those that express Arginase 1 (Arg) can be considered monocytic myeloid-derived suppressor cells (mMDSCs). IACS-70654 reduced mMDSC infiltration in 2208L and PyMT-N tumors (Supplemental Figure S2F). These results suggested that IACS-70654 can reduce immunosuppression and might be an effective therapy for neutrophil-enriched TNBC, which has been correlated with poor patient outcomes and therapy resistance (Kim et al., 2019, Wu and Zhang, 2020). Because IACS-70654 reduced neutrophils in tumors and blood, we further hypothesized that IACS-70654 might inhibit the abnormal neutrophil generation in the bone marrow promoted by tumor outgrowth (Casbon et al., 2015, Hao et al., 2023, Veglia et al., 2018).

### IACS-70654 reprogrammed and reduced the proliferation of neutrophils in the bone marrow

To characterize the changes in bone marrow after IACS-70654 treatment, we first performed scRNA-seq analyses on all CD45-positive cells collected from the bone marrow of non-tumor-bearing WT BALB/c and 2208L tumor-bearing mice. The 2208L tumor-bearing mice were treated with vehicle or IACS-70654 for 6 days (5 total treatments), and bone marrow was collected 24 hrs after the last treatment. As expected, the fraction of neutrophils was increased in 2208L tumors with a concomitant decrease in the fraction of monocytes and dendritic cell progenitors (pDCs) (**Figure 3A and B** and Supplemental Figure S3A). IACS-70654 treatment reduced the production of bone marrow neutrophils and restored that of monocytes and pDCs (**Figure 3B**). This observation was also confirmed by flow cytometry analyses coupled with measuring BrdU incorporation (**Figure 3C** and Supplemental Figure S3B). The neutrophils then were isolated and clustered to identify subpopulations using previously identified markers (Carnevale et al., 2023, Grieshaber-Bouyer et al., 2021, Qu et al., 2023) (**Figure 3D**). Pro-neutrophils (proNeu) and Pre-neutrophils (preNeu) are the proliferative precursors (Qu et al., 2023). Therefore, we examined the cell cycle score of the proliferative proNeu and preNeu and found that IACS-70654 decreased the fraction of cells exhibiting a G2/M signature (**Figure 3E**). Moreover, IACS-70654 reduced the expression of *Csf3r,* which encodes the granulocyte colony stimulating factor receptor, the receptor for the granulocyte colony stimulating factor (G-CSF), in mature neutrophils (**Figure 3F**). G-CSF level was upregulated by the 2208L tumor (Supplemental Figure S3C). IACS-70654 might reduce G-CSF signaling, a critical driver for neutrophil production, migration, and immunosuppression (Casbon et al., 2015, Karagiannidis et al., 2021, Waight et al., 2011). IACS-70654 also decreased the RNA expression of C-C chemokine receptor type 1 (CCR1) and Ly6G, both of which are critical for neutrophil migration (Wang et al., 2012, Metzemaekers et al., 2020) (**Figure 3F, G** and Supplemental Figure S3D). In addition, we discovered that bone marrow neutrophils in 2208L tumor-bearing mice downregulated the expression of *Thbs1*, which encodes Thrombospondin-1 (TSP-1) (**Figure 3I** and Supplemental Figure S3E). IACS-70654 restored the RNA expression of TSP-1 (**Figure 3I** and Supplemental Figure S3D). Moreover, cystatins (*Stfa2*, *Stfa3, Stfa2l1, Cstdc4, Cstdc5, Cstdc6*) expression was significantly upregulated in immature and mature neutrophils after IACS-70654 treatment (Supplemental Figure S3D-G). TSP-1 and cystatins are both inhibitors of neutrophil serine protease (Pham, 2006, Zhao et al., 2015). Neutrophil serine proteases were reported to promote neutrophil release into the blood by reducing C-X-C motif chemokine ligand 12 binding to C-X-C chemokine receptor type 4. They can also be involved in the protumor activity of neutrophils by inducing the release of C-X-C motif chemokine ligand 2, which promotes neutrophil recruitment, and activates IL-1β, a protumor cytokine (Baker et al., 2019, Cambier et al., 2023). These results revealed that IACS-70654 might reduce the proliferation, migration, and protumor activity of bone marrow neutrophils. However, hematopoietic stem and progenitor cells (HSPCs) may also be affected by tumor-derived factors (Hao et al., 2023, Casbon et al., 2015). Accordingly, we next investigated the changes in HSPCs after IACS-70654 treatment.

**Figure 3.**
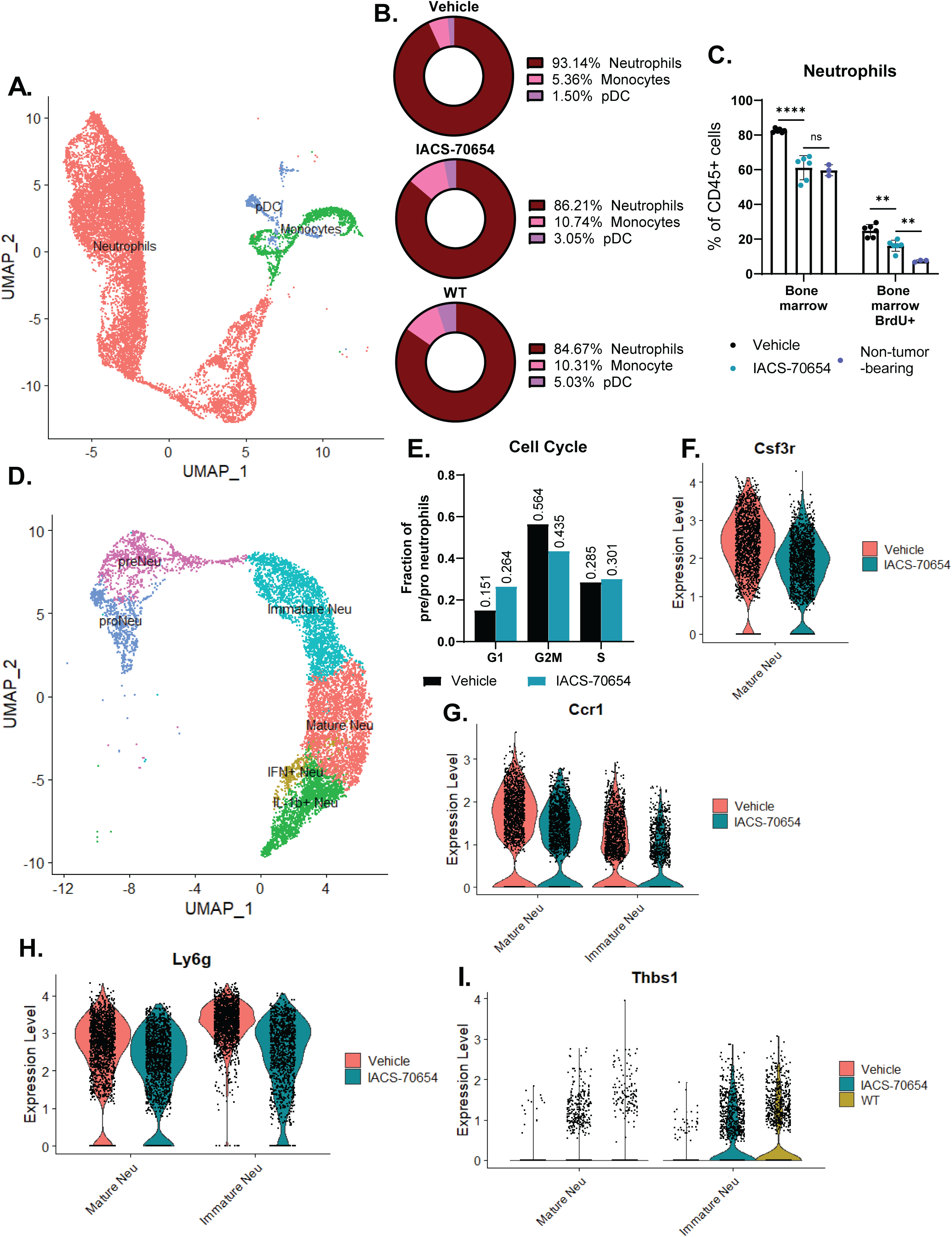
IACS-70654 reprogrammed bone marrow neutrophils. **A.** UMAP plot of bone marrow myeloid cells (neutrophils, monocytes, and pDCs) with annotations. **B.** Fractions of bone marrow myeloid cells in 2208L tumor-bearing (treated with vehicle or IACS-70654) and untreated non-tumor-bearing WT BALB/c mice from scRNA-seq analyses. **C.** Flow cytometry analyses of neutrophils in the bone marrow of 2208L tumor-bearing mice treated with vehicle or IACS-70654 and untreated non-tumor-bearing WT BALB/c mice. BrdU was administered 24 hrs before the analysis. Ordinary one-way ANOVA and Tukey’s multiple comparisons test were used. Five biological replicates were used for each treatment arm of 2208L tumor-bearing mice, and 3 biological replicates were used for non-tumor-bearing mice. ****, p<0.0001; **, p<0.01; ns, p>0.05. Error bars represent SD. **D.** UMAP plot of bone marrow neutrophils with annotations. **E.** Fractions of pre-neutrophil and pro-neutrophil in G1, G2M, or S cell cycle stage in the bone marrow of 2208L tumor-bearing mice treated with vehicle or IACS-70654 from scRNA-seq analyses. **F.** Violin plot showing the expression level of *Csf3r* in mature neutrophils in the bone marrow of 2208L tumor-bearing mice treated with vehicle or IACS-70654. Adjusted p = 3.29e-92. **G.** Expression level of *Ccr1* in mature and immature neutrophils in the bone marrow of 2208L tumor-bearing mice treated with vehicle or IACS-70654. Mature Neutrophils: adjusted p = 9.39e-46; Immature neutrophils: adjusted p = 5.95e-39. **H.** Violin plot showing the expression level of *Ly6g* in immature and mature neutrophils of 2208L tumor-bearing mice treated with vehicle or IACS-70654. **I.** Expression levels of *Thbs1* in mature and immature neutrophils in the bone marrow of 2208L tumor-bearing (treated with vehicle or IACS-70654) and non-tumor-bearing WT BALB/c mice.

### IACS-70654 induced transcriptional changes in HSPCs to reduce abnormal myelopoiesis

To determine whether IACS-70654 can reprogram the abnormal myelopoiesis induced by the neutrophil-enriched tumor, we performed scRNA-seq on HSPCs (defined as CD45^+^Lin^-^C-kit^+^) in the bone marrow of 2208L tumor-bearing mice treated with vehicle or IACS-70654 and non-tumor-bearing mice. The dataset was filtered to contain only HSPCs involved in myelopoiesis and annotated using classic HSPC markers as described previously (Hao et al., 2023, Yanez et al., 2017). Two distinct populations of granulocyte-monocyte progenitors (GMPs) and common myeloid progenitors (CMPs) were found (**Figure 4A**). Based on clustering and trajectory analyses, GMP-1 and CMP-1 were determined to be involved in myelopoiesis (**Figure 4A**, Supplemental Figure S4A). 2208L tumors upregulated the fraction of GMP-1, monocyte progenitors (MPs), and proNeu, whereas IACS-70654 treatment reduced those progenitors, indicating an inhibition of myelopoiesis (**Figure 4B**). In cluster 4, a CMP-1 cluster, IACS-70654 led to the downregulation of myeloid differentiation and activation pathways (**Figure 4C**). In addition, in CMP-1 IACS-70654 decreased the expression of *Prtn3* and *Ms4a3*, which have been associated with the differentiation and proliferation of myeloid progenitor cells (Sköld et al., 1999, Ishibashi et al., 2018) (**Figure 4D and E**). Moreover, IACS-70654 decreased the expression of genes that were correlated with the differentiation of early progenitors to neutrophils (*Ifitm1, Ifitm3, Calr, Cd63, Plac8, Prtn3, Ms4a3, Igfbp4*) in CMP-1 and multipotent progenitors (MPPs) (**Figure 4D and E**, Supplemental Figure S4B-F). These changes imply that IACS-70654 might reduce the differentiation of HSPCs to neutrophils. Moreover, in CMP-1, MMPs, and hematopoietic stem cells (HSCs), IACS-70654 induced the expression of *Malat1*, a long non-coding RNA shown to inhibit differentiation of early HSPCs (Ma et al., 2015) (**Figure 4F**). In HSCs and MPPs, IACS-70654 also increased the expression of *Txnip*, which can play a role in keeping HSCs in an undifferentiated state and reducing their mobility (Jeong et al., 2009) (Supplemental Figure S4C and D). Furthermore, in the peripheral blood of 2208L tumor-bearing mice, IACS-70654 reduced the level of IL-3, a cytokine known to induce HSC and myeloid differentiation (Nitsche et al., 2003, Johnson et al., 2002) (Supplemental Figure S4G). These observations suggest that IACS-70654 treatment might retain the early HSPCs in an undifferentiated state, explaining the accumulation of HSC, MPP, and CMP-1 (**Figure 4B**).

**Figure 4.**
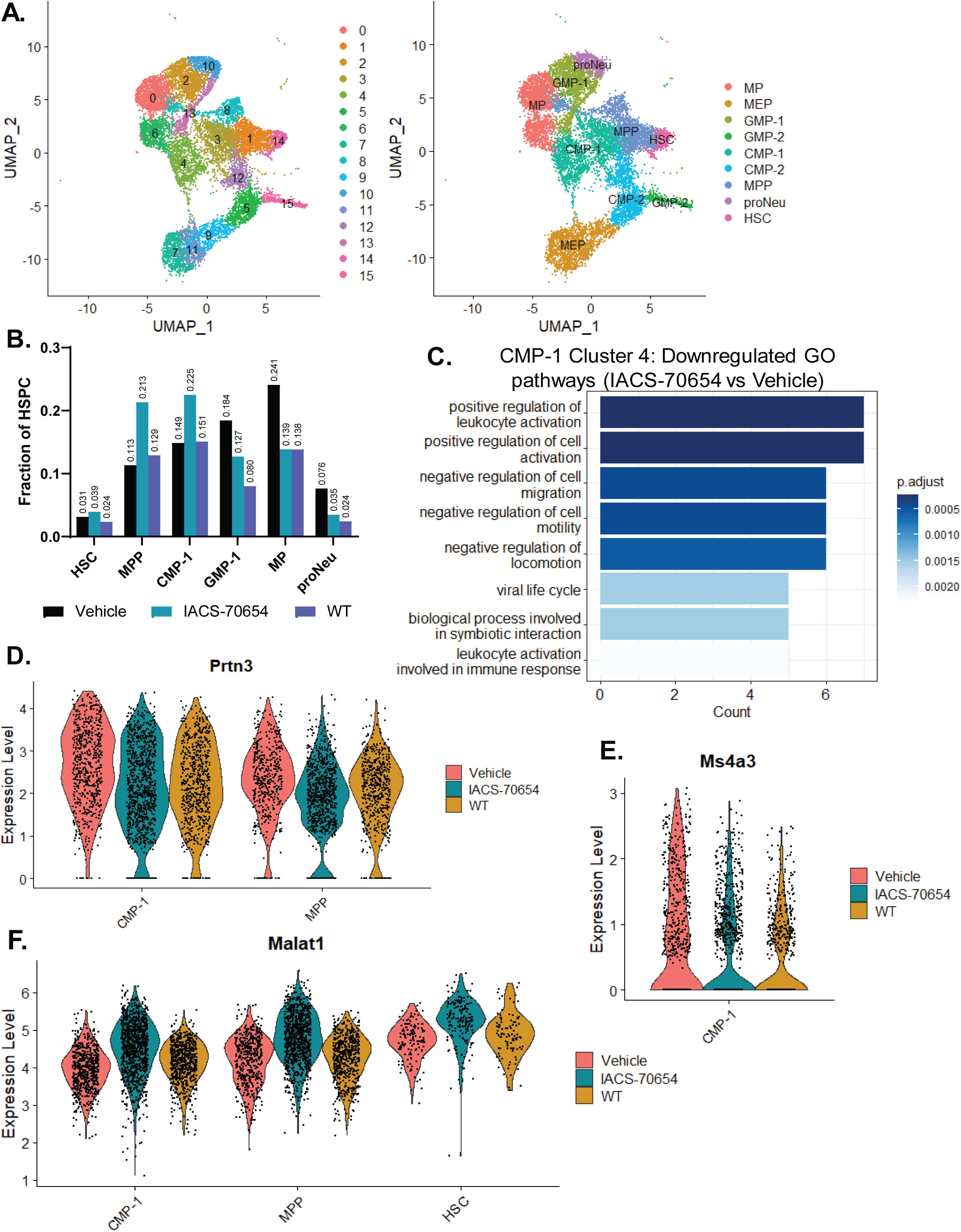
IACS-70654 reprogrammed abnormal myelopoiesis induced by the neutrophil-enriched tumor. **A.** UMAP plot of HSPC clusters without (left) and with (right) annotation. **B.** Fraction of myeloid progenitor cells in HSPCs of 2208L-tumor bearing mice (treated with vehicle or IACS-70654) and non-tumor-bearing WT mice. **C.** GO pathway enrichment analysis of the significantly downregulated genes (Log_2_ fold change < -0.5 and adjusted p-value < 0.01) in cluster 4 of HSPCs of 2208L tumor-bearing mice treated with IACS-70654 versus vehicle. Biological Process gene sets from the GO database were used. The top 8 pathways were listed with numbers of genes enriched. **D.** Violin plots showing the RNA expression of *Prtn3* in CMP-1 and MPPs of 2208L tumor-bearing mice treated with vehicle or IACS-70654 and non-tumor-bearing WT mice. **E.** Violin plot showing the RNA expression of *Ms4a3* in CMP-1 of 2208L tumor-bearing mice treated with vehicle or IACS-70654 and non-tumor-bearing WT mice. **F.** Violin plots showing the RNA expression of *Malat1* in CMP-1, MPPs, and HSCs of 2208L tumor-bearing mice treated with vehicle or IACS-70654 and non-tumor-bearing WT mice.

Taken together, these results suggested that IACS-70654 reduced the differentiation of early HPSCs to reduce the overproduction of GMP-1 and thus neutrophils. Besides neutrophils, IACS-70654 may also elicit effects on 2208L tumor cells. Thus, we next investigated the effects of IACS-70654 in tumor cells.

### IACS-70654 induced both an IFN response and antigen presentation in tumor cells

To investigate how IACS-70654 impacts tumor cells, we again utilized the scRNA-seq dataset from 2208L tumors treated with vehicle or IACS-70654 for 7 days and analyzed the tumor cell cluster (Supplemental Figure S5A and B). IACS-70654 induced the expression of MHCI components (*H2-D1, B2m, H2-K1, H2-Q4, H2-Q7*) (**Figure 5A-C**). Using flow cytometry, we confirmed that IACS-70654 treatment induced the protein expression of MHCI on the cell surface of 2208L tumor cells *in vivo* (**Figure 5D**). Besides MHCI, IACS-70654 also induced the expression of genes associated with IFNβ-and virus-response pathways (**Figure 5A, B and C**). Using a cytokine/chemokine array, we detected an elevated level of IFNβ in 2208L tumors treated with IACS-70654 as compared to controls treated with vehicle (**Figure 5E**). IACS-70654 also downregulated genes involved in the regulation of inflammation and inhibition of cytokine production, such as *Cd200* and *Cebpb* which have been shown to induce immunosuppression in the TIME (Choe and Choi, 2023, Matherne et al., 2023) (**Figure 5A** and Supplemental S5C). In tumor cells, these results showed that in 2208L tumor cells IACS-70654 induced both MHCI expression and an IFN response, suggesting induced antigen presentation and immune stimulation including T-cell activation. Therefore, we next investigated the effects of IACS-70654 on tumor-infiltrated lymphocytes and the response of neutrophil-enriched tumors to ICB.

**Figure 5.**
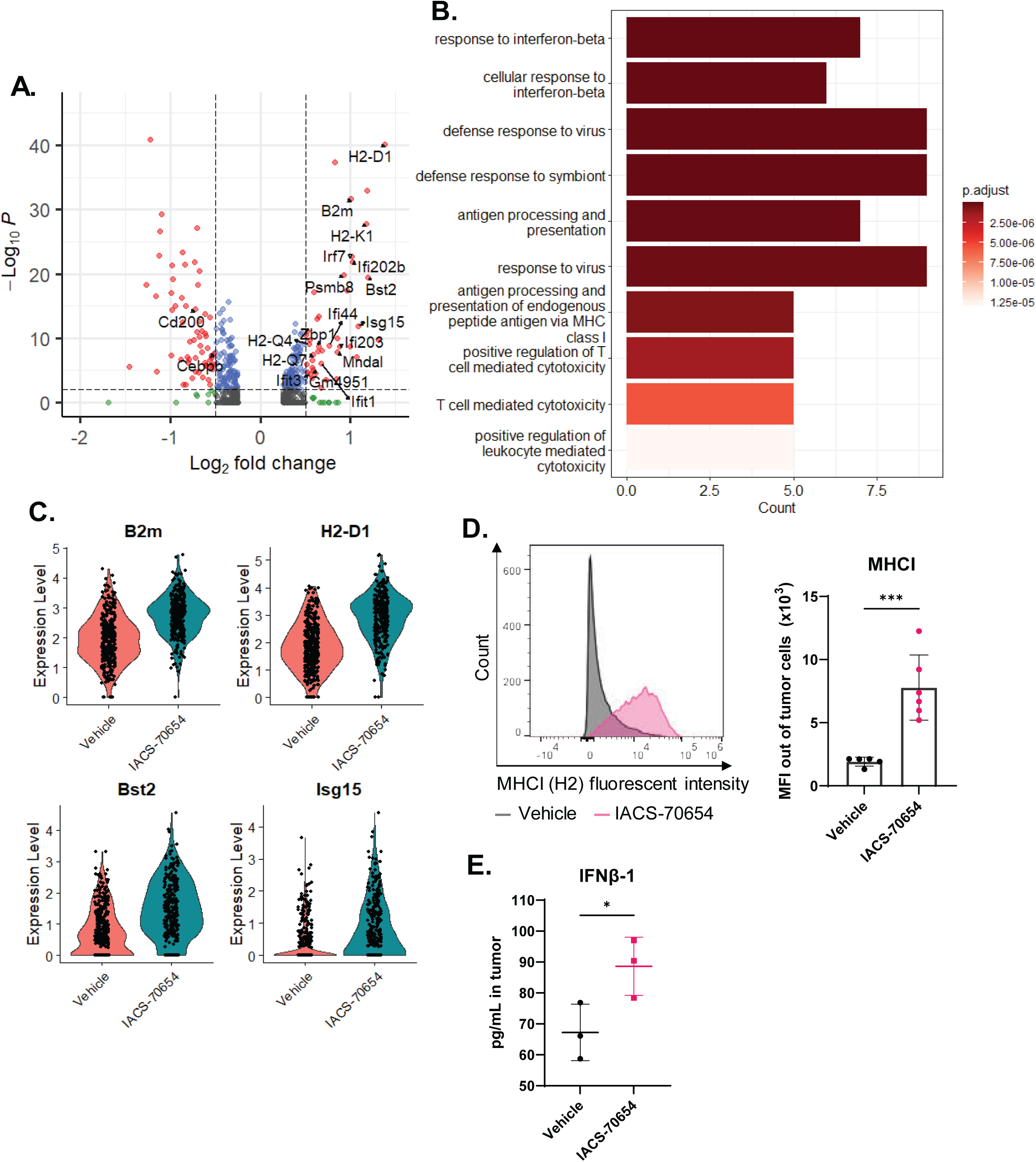
IACS-70654 induced an IFN-associated response and MHC I expression in 2208L tumor cells. **A.** Volcano plot showing –Log_10_ p-value versus Log_2_ fold change in RNA expressions in 2208L tumor cells treated *in vivo* with IACS-70654 versus those treated with vehicle. Genes that showed significant changes (Log_2_ fold change > 0.5 or < -0.5 and adjusted p-value < 0.01) in expression are represented by red dots. The genes that are associated with IFN response, antigen representation, and immunosuppression are labeled. **B.** GO pathway enrichment analysis of the significantly upregulated genes (Log_2_ fold change > 0.5 and adjusted p-value < 0.01) in 2208L tumor cells treated *in vivo* with IACS-70654 versus those treated with vehicle. Biological Process (BP) gene sets from the GO database were used. The top 10 pathways were listed with numbers of genes enriched. **C.** Violin plots showing RNA expression of representative MHCI components (*B2m* and *H2-D1*) and IFN response genes (*Bst2* and *Isg15*) in 2208L tumor cells treated *in vivo* with vehicle or IACS-70654. **D.** Flow cytometry analysis of MHCI expression in 2208L tumor cells treated *in vivo* with vehicle or IACS-70654. Left: Representative histograms of MHCI expression in 2208L tumor cells (CD45-/TER119-/CD31-/EpCAM+). Right: Median fluorescent intensity (MFI) of MHCI in 2208L tumor cells. For the vehicle-treated group, five biological replicates were used, and for the IACS-70654-treated group, six biological replicates were used. Two-tailed unpaired Student’s *t* test was used. ***, p<0.001. Error bars represent SD. **E.** Quantification of IFNβ level in tissue homogenate from 2208L tumors treated with IACS-70654 or vehicle by the cytokine/chemokine array. Three biological replicates were used for each treatment arm. Two-tailed unpaired Student’s *t* test was used. Error bars represent standard deviation. *, p<0.05.

### IACS-70654 activated CTLs and improved response to ICB

Neutrophil-enriched TNBC models such as the 2208L model usually have very low lymphocyte infiltration and complete resistance to ICB (Kim et al., 2019). After a 7-day treatment, 2208L tumors treated with IACS-70654 showed a significantly higher level of CTL (CD3^+^/CD8^+^) infiltration (**Figure 6A**). This finding was confirmed using immunostaining, and CTLs were observed in the tumor center instead of stroma after IACS-70654 treatment (**Figure 6B**). To determine whether CTLs play a critical role in tumor growth inhibition by IACS-70654, 2208L tumor-bearing mice were treated with an anti-CD8 antibody. The anti-CD8 antibody was administered 24 hrs before starting IACS-70654 treatment and then throughout the experiment.

**Figure 6.**
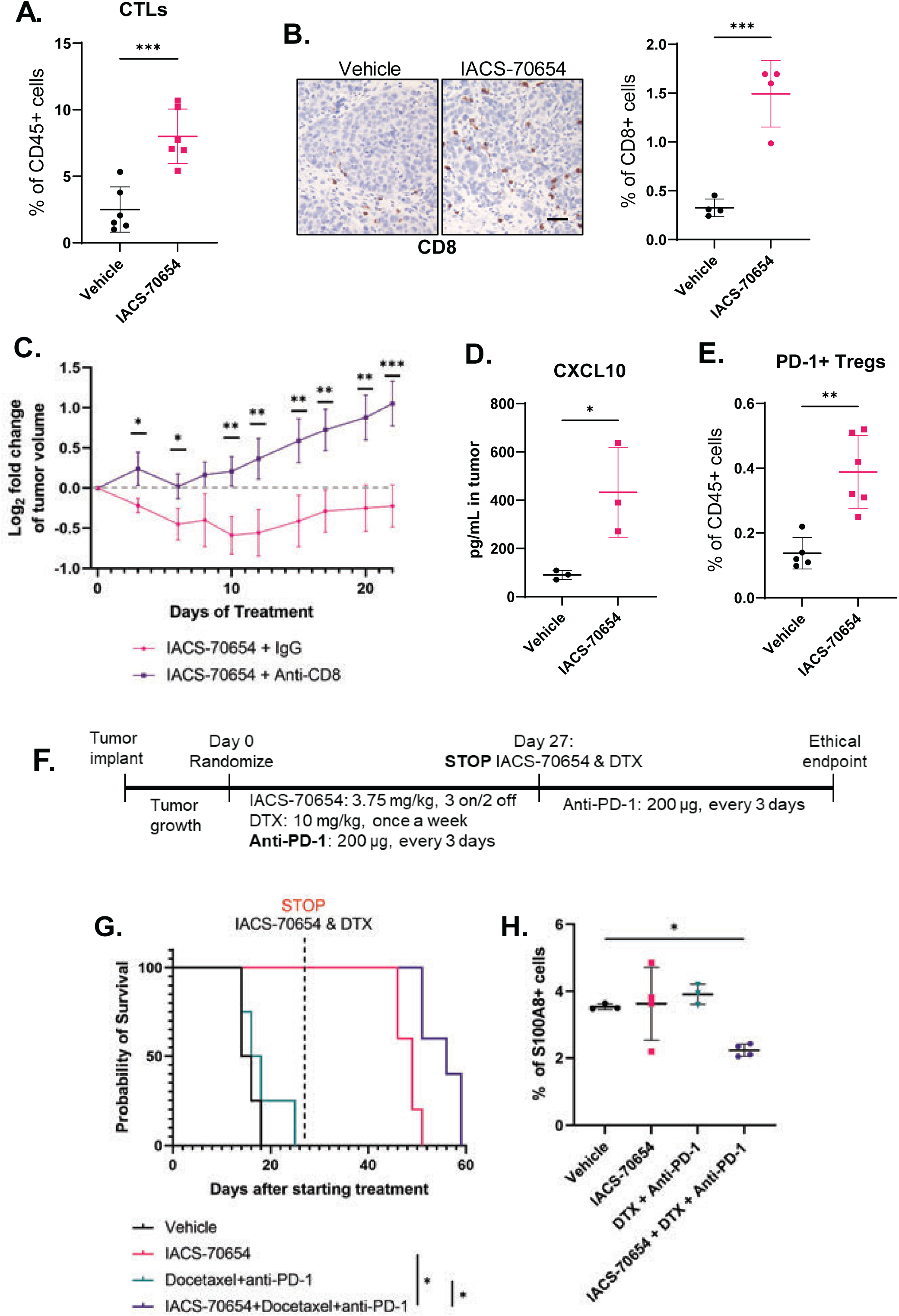
IACS-70654 induced a CTL-dependent response and improved response to ICB. **A.** Flow cytometry analysis of CTLs in the 2208L tumors treated with vehicle or IACS-70654 for 7 days. Six biological replicates were used for each treatment arm. Two-tailed unpaired Student’s *t* test was used. ***, p<0.001. Error bars represent SD. **B.** Left: Representative images of immunostaining of CD8 on sections of 2208L tumors treated with vehicle or IACS-70654 for 7 days. Scale bar: 50 µm. Right: Quantification of CD8 staining using Fiji. Three representative images were analyzed for each tumor, and four biological replicates were used for each treatment arm. Two-tailed unpaired Student’s *t* test was used. ***, p<0.001. Error bars represent SD. **C.** Log_2_ fold change of 2208L tumor volume with CTL depletion. 2208L tumors were treated with Anti-CD8 24 hrs before starting IACS-70654 treatment. For each treatment arm, *n* =5. Two-way ANOVA and Sidak’s multiple comparisons test were used. *, p<0.05; **, p<0.01; ***, p<0.001. Error bars represent SD. **D.** Quantification of CXCL10 level in tissue homogenate of 2208L tumors treated with IACS-70654 or vehicle by the cytokine/chemokine array. Three biological replicates were used for each treatment arm. Two-tailed unpaired Student’s *t* test was used. Error bars represent standard deviation. *, p<0.05. **E.** Flow cytometry analysis of infiltrated PD-1+ regulatory T cells (Tregs) in the 2208L tumors treated with vehicle or IACS-70654 for 18 days. For each group, five biological replicates were used. Two-tailed unpaired Student’s *t* test was used. ***, p<0.001. Error bars represent SD. **F.** Treatment schemes of IACS-70654 in combination with anti-PD-1 and DTX. Tumor implant, growth, and randomization were performed as previously described. DTX was administered i.p. weekly at 10mg/kg, half of the clinically equivalent dose. Anti-PD-1 was administered i.p. every three days at 200 µg per mouse. On Day 27 after starting treatment, IACS-70654 and DTX treatments were stopped, and anti-PD-1 treatment continued until all tumors reached the ethical endpoint (tumor size ≥1,500mm^3^). **G.** Kaplan-Meier survival curves of 2208L tumors that reached the ethical endpoint in the combination treatment study. For vehicle and DTX + anti-PD-1 treated groups, *n* = 4. For IACS-70654 and combination-treated groups, *n* = 5. Log-rank test with Bonferroni correction was used. *, p<0.05. **H.** Quantification of the immunostaining of S100A8 in 2208L tumors from all groups. Ordinary one-way ANOVA and Tukey’s multiple comparisons test were used. Three biological replicates were used for vehicle and DTX + anti-PD-1 treated groups. Four biological replicates were used for IACS-70654 and combination-treated groups. *, p>0.05. Error bars represent SD.

Successful depletion of CTLs in the tumors was confirmed by flow cytometry (Supplemental Figure S6A). CTL depletion attenuated the effects of IACS-70654 and suggested that CTLs were important in both the immediate tumor regression and the durable inhibition of tumor growth (**Figure 6C**). Moreover, in 2208L tumors, IACS-70654 induced the level of C-X-C motif chemokine ligand 10 (CXCL10), a chemokine contributing to CTL recruitment and associated with better efficacy of ICB (**Figure 6D**) (Reschke and Gajewski, 2022). IACS-70654 did not increase programmed cell death protein 1 (PD-1)-positive CTLs but increased PD-1+ regulatory T cells (Tregs, defined as CD4^+^FoxP3^+^), implying a potential benefit of combining IACS-70654 with anti-PD-1 (**Figure 6E** and Supplemental Figure S6B). Thus, we tested the efficacy of IACS-70654 in combination with docetaxel (DTX) and anti-PD-1, which partially mimics the current standard-of-care therapy for TNBC (**Figure 6F**). DTX is known to cause adverse effects in breast cancer patients, and lowering the dose is commonly used to mitigate adverse effects (Ho and Mackey, 2014, Loeser et al., 2024). Therefore, in this study, DTX was administered at 10 mg/kg, which is half of the clinically relevant dose. The combination treatment was well tolerated and showed no signs of toxicity (Supplemental Figure S6C). DTX in combination with anti-PD-1 did not significantly improve the survival of 2208L tumor-bearing mice and failed to inhibit tumor growth (**Figure 6G** and Supplemental Figure S6D). All tumors treated with vehicle or DTX in combination with anti-PD-1 reached the ethical endpoint within 25 days. IACS-70654 in combination with DTX and anti-PD-1, similar to IACS-70654 alone, durably inhibited the tumor growth (Supplemental S6D). To determine whether the combination treatment has durable long-term effects on the response of 2208L tumors to anti-PD-1, we stopped IACS-70654 and DTX treatment in the remaining 2 groups on Day 27 (**Figure 6G**). Compared to IACS-70654 alone, IACS-70654 in combination with DTX and anti-PD-1 significantly delayed the regrowth of 2208L tumors (**Figure 6G**). Furthermore, immunostaining of the tumors at the endpoint revealed that TANs again accumulated in the tumor treated with IACS-70654, indicating immune suppression (**Figure 6H**). However, tumors treated with IACS-70654 in combination with DTX and anti-PD-1 were infiltrated with significantly fewer TANs, which might explain the delayed recurrence (**Figure 6H**). In summary, IACS-70654 inhibited the growth of 2208L tumors in a CTL-dependent manner and can potentially improve the response of neutrophil-enriched TNBC to standard-of-care therapies. Although we demonstrated the efficacy of IACS-70654 treatment in primary tumors of neutrophil-enriched TNBC models, the clinically unmet need is to treat metastasis. Accordingly, we next investigated the effects of IACS-70654 in the metastatic setting.

### IACS-70654 inhibited the growth of established lung metastases

From the scRNA-seq analyses of 2208L primary tumors treated with IACS-70654, we observed that IACS-70654 downregulated the expression of genes associated with migration and negative regulation of cell adhesion pathways (Supplemental Figure S5C). More importantly, many of the downregulated genes (*Fn1, Hspb1, Postn, Mia, Fgfr1, Serpine2)* have been associated with tumor metastasis (Glasner et al., 2018, Guba et al., 2000, Xie et al., 2020, Huo et al., 2023, Labreche et al., 2021, Smirnova et al., 2016) (**Figure 7A** and Supplemental Figure S7A). We also observed a reduction in fibroblast growth factor receptor 1 (FGFR1) protein expression in 2208L primary tumors treated with IACS-70654 (Supplemental Figure S7B). Next, we used an experimental metastasis model to test the effects of IACS-70654 on established lung metastases in the 2208L model. To generate lung metastases, 100,000 cells dissociated from a 2208L tumor were injected into the tail vein of each WT BALB/c mouse (**Figure 7B**).

**Figure 7.**
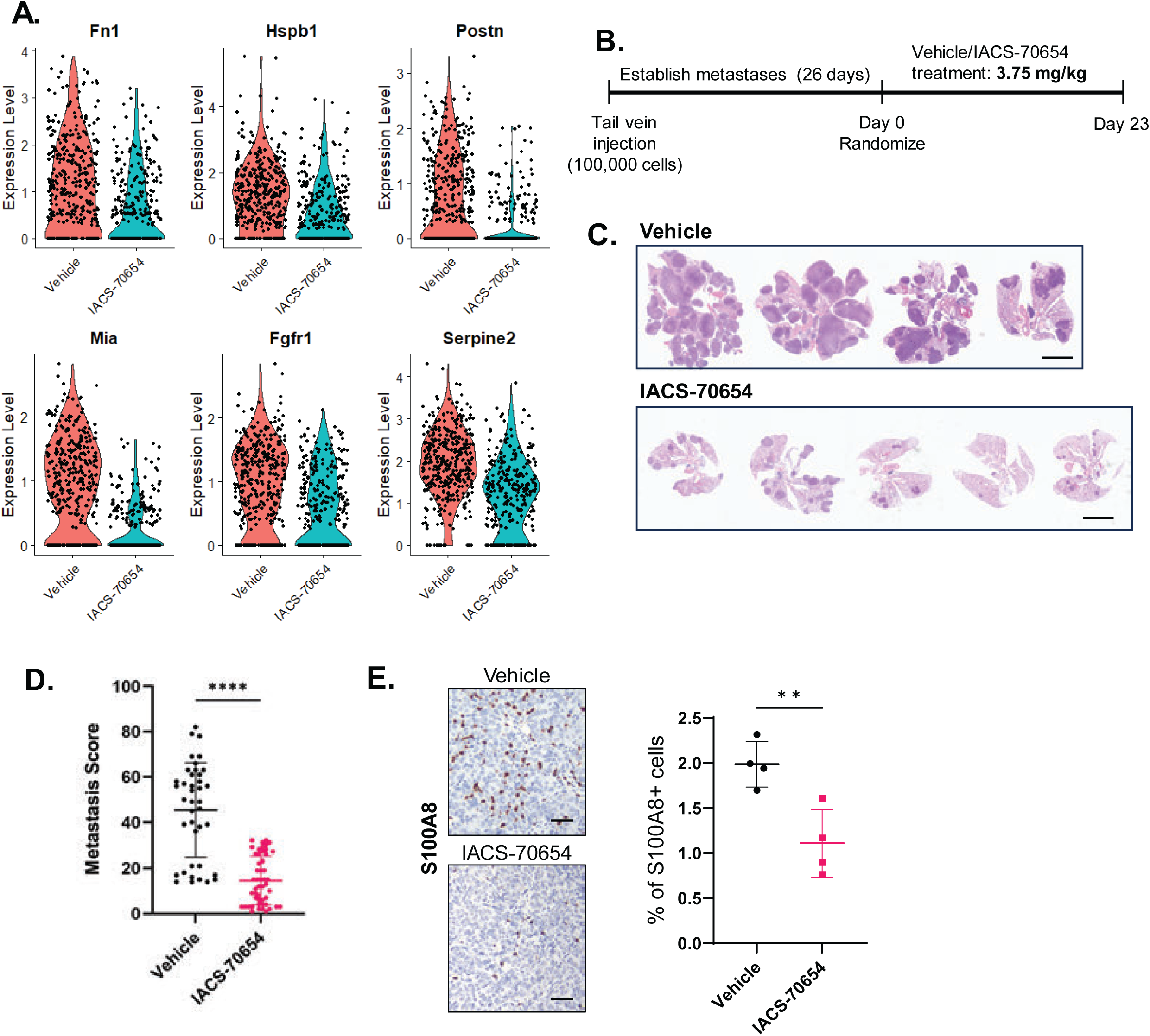
IACS-70654 reduced the growth of established 2208L lung metastases. **A.** Violin plots showing RNA expression of representative genes that might be involved in migration and metastasis in 2208L tumor cells treated with vehicle and IACS-70654. Adjusted p<0.01 for all genes. **B.** Experimental design for studying the effects of IACS-70654 on established lung metastases. 100,000 dissociated 2208L tumor cells were injected into the tail vein of each wild-type BALB/c mouse. On Day 0 of the treatment, the mice were randomized, and then the treatment was initiated. Vehicle or IACS-70654 at 3.75 mg/kg was administered orally on a 3-on/2-off schedule. The mice were treated for 23 days when all vehicle-treated mice reached the ethical endpoint. One mouse from the Vehicle group had to be euthanized on Day 22 due to poor body condition. **C.** Representative images from H&E staining of lungs with 2208L metastases from mice treated with vehicle or IACS-70654. Scale bar: 5 mm. **C.** Quantifications of metastases in the lungs of mice treated with vehicle or IACS-70654. Serial sectioning was performed to collect a total of 10 sections for each sample. H&E staining was performed on all sections. The metastases were categorized and counted based on the diameter. Metastases with sizes of <1 mm, 1-3 mm, 3-5 mm, and >5 mm were assigned scores of 1, 2, 3 and 4, respectively. For the vehicle group, *n* = 4, and for the IACS-70654 group, *n* = 5. Two-tailed unpaired Student’s *t* test was used. ****, p<0.0001. Error bars represent SD. **E.** Left: Representative images of immunostaining of S100A8 on 2208L lung metastasis sections treated with vehicle or IACS-70654. Scale bar: 50 µm. Right: Quantification of S100A8 staining using Fiji. Three representative images were analyzed for each tumor, and four biological replicates were used for each treatment arm. Two-tailed unpaired Student’s *t* test was used. **, p<0.01. Error bars represent SD.

Because results from the primary tumor indicated that IACS-70654 mainly affected the immune response, unlabeled tumor cells were injected to prevent the introduction of neoantigens derived from fluorescent reporters (Grzelak et al., 2022). We collected the lungs from two mice before starting treatment to ensure the successful establishment of lung metastases. The mice were randomized and treated with vehicle or IACS-70654 for 23 days until most vehicle-treated mice reached the ethical endpoint (**Figure 7B**). From H&E staining, we observed that the lung metastatic burden of IACS-70654-treated mice was decreased as compared to vehicle-treated mice (**Figure 7C**). The metastatic burden was quantified by counting the metastases using serial sectioning and a size-based scoring system to calculate a metastasis score for each lung section (**Figure 7D**). The metastasis score of IACS-70654-treated mice was significantly lower than that of vehicle-treated mice (**Figure 7D**). In the plasma of 2208L lung metastasis-bearing mice, we observed an abnormal upregulation of chemokine ligand 19 (CCL19), which has been associated with breast cancer metastasis (Gowhari Shabgah et al., 2022) (Supplemental Figure S7C). The CCL19 level in the plasma decreased with IACS-70654 treatment (Supplemental Figure S7C). Moreover, neutrophils have been shown to promote metastasis, and 2208L retained its neutrophil-enriched signature in the lung (Wu and Zhang, 2020) (**Figure 7E**). IACS-70654 reduced the number of infiltrated neutrophils in 2208L lung metastases (**Figure 7E**). IACS-70654 also reduced Ly6G expression in the circulating neutrophils of 2208L lung metastasis-bearing mice (Supplemental Figure S7D). Thus, since IACS-70654 inhibited established lung metastasis in these preclinical models it may potentially provide a therapeutic alternative for the treatment of metastatic neutrophil-enriched TNBC.

## Discussion

This study investigated the effects of CBP/P300 BRD inhibition on four syngeneic preclinical models of TNBC using IACS-70654, a novel and selective inhibitor of CBP/P300 BRD. It demonstrated that IACS-70654 inhibited the growth of neutrophil-enriched TNBC models in part by reducing immunosuppressive TANs and activating immune responses.

CBP/P300 BRD inhibition has been reported to reprogram cancer cells to reduce proliferation and treatment resistance in many cancer types, but most studies and clinical trials have focused on blood cancers and prostate cancer (Chen et al., 2022, Nicosia et al., 2023, Raisner et al., 2018, Jin et al., 2017). For TNBC, CBP/P300 BRD inhibition has been shown to reduce the growth of patient-derived androgen receptor-positive TNBC xenografts and MDA-MB-231 xenografts (Caligiuri et al., 2023, de Almeida Nagata et al., 2019). CBP/P300 BRD inhibitors have not been tested in any immunocompetent models of TNBC, and therefore our study reports we believe for the first time the effects of a CBP/P300 BRD inhibitor in syngeneic TNBC models. We confirmed that CBP/P300 BRD inhibition can reduce or inhibit tumor proliferation in several neutrophil-enriched TNBC models, and our findings suggested that high TAN frequency might predict a better response of TNBC to CBP/P300 BRD inhibitors.

Investigation of the TIME after CBP/P300 BRD inhibition revealed a substantial reduction of TANs in most models tested regardless of TAN frequencies. TANs in neutrophil-enriched models were shown to be immunosuppressive and considered MDSCs, suggesting that the growth of neutrophil-enriched models might be inhibited after TAN reduction (Kim et al., 2019). The effects of CBP/P300 BRD inhibition on MDSCs of TNBC have previously been studied using an MDA-MB-231 xenograft model in immunocompromised hosts, and our observations are in general consistent with what was previously reported (de Almeida Nagata et al., 2019). However, MDA-MB-231 is a mesenchymal TNBC model, in which TAMs but not TANs were the most abundant myeloid cells, and the macrophage-enriched mesenchymal T12 tumors did not respond significantly to CBP/P300 BRD inhibition. (de Almeida Nagata et al., 2019). This difference may be explained by the nature of a human tumor xenograft model. Since all the stromal cells including TAMs are of murine origin, they might lead to weaker support for tumor growth and possibly less immunosuppressive TIME (Morgan, 2012). Moreover, CBP/P300 BRD inhibition certainly does not affect only TANs. In agreement with the previous study, we also observed reduced mMDSCs by CBP/P300 BRD inhibition (de Almeida Nagata et al., 2019). In addition, we observed an increase in the fraction of a TAM subtype expressing an IFN response signature which might contribute to immune stimulation and tumor inhibition. Therefore, those changes might be sufficient to reduce tumor growth significantly in a macrophage-enriched xenograft model but not in a syngeneic model. Overall, our study suggested that CBP/P300 BRD inhibition reduces immunosuppression primarily by reducing TANs, and therefore neutrophil-enriched TNBC may be more sensitive to CBP/P300 BRD inhibition. Furthermore, we have previously observed that several macrophage-enriched TNBC models initially responded to ICB, but rapidly recurred concomitant with the appearance of immunosuppressive TANs (5). In several of these models, there appears to be a Yin Yang relationship between immunosuppressive TAMs and TANs. This may provide an escape mechanism for TNBC treated with ICB and suggests that targeting TANs by CBP/P300 BRD inhibition may provide an important therapeutic option since these neutrophil-enriched tumors are in general resistant to ICB.

Previous studies have reported that bone marrow is the primary site responsible for abnormal neutrophil production induced by neutrophil-enriched tumors. Bone marrow neutrophils from 2208L and PyMT-N tumor-bearing mice were shown to suppress T-cell activation *in vivo* (Casbon et al., 2015, Kim et al., 2019). CBP/P300 BRD inhibition has been shown to reprogram acute myeloid leukemia and myeloma cells in the bone marrow (Nicosia et al., 2023). Abnormal proliferation is a shared characteristic between hematological malignancies and neutrophil overproduction caused by neutrophil-enriched TNBC. In this study, we confirmed the overproduction of neutrophils in the bone marrow of 2208L tumor-bearing mice and showed that IACS-70654 reduced the proliferation of neutrophil precursors. The current study also indicated that CBP/P300 BRD inhibition may disrupt the release of neutrophils into the bloodstream. Then we investigated the early progenitors and determined that the presence of 2208L tumors increased both HSCs and GMP-1, the GMP population shown to differentiate towards neutrophils. These findings agreed with both flow cytometry and scRNA-seq results from a recent study from our group, which also used multiple neutrophil-enriched models including 2208L and PyMT-N models (Hao et al., 2023). This recent study also demonstrated that the genes associated with the differentiation and proliferation of myeloid progenitors were upregulated while those associated with the inhibition of myeloid differentiation were downregulated (Hao et al., 2023). In the present study, we found that CBP/P300 BRD inhibition reduced GMP-1 and reversed the alternations of gene expression induced by the neutrophil-enriched tumor. Furthermore, our results suggested that CBP/P300 BRD inhibition downregulated the differentiation of early progenitors including CMPs, MPPs, and possibly HSCs, trapping the progenitor cells in an undifferentiated state and leading to reduced GMP-1. Taken together, these results suggest that CBP/P300 BRD inhibition reprogrammed the differentiation and inhibited the proliferation of neutrophil progenitors in the bone marrow to reduce neutrophil overproduction. These findings again emphasize the potential therapeutic benefits of CBP/P300 BRD inhibitors on neutrophil-enriched TNBC.

Besides TAN reduction, CBP/P300 BRD inhibition was demonstrated to stimulate the immune response. MHCI expression is downregulated in many cancer types to promote immune evasion and low MHCI expression has been associated with poor response to ICB (Taylor and Balko, 2022, Dhatchinamoorthy et al., 2021). MHCI expression can be silenced epigenetically, and thus epigenetic inhibitors have been suggested as one potential treatment that might restore MHCI expression (Taylor and Balko, 2022). CBP/P300 BRD inhibition induced MHCI expression in 2208L tumor cells *in vivo*, suggesting that CBP/P300 BRD might be involved in MHCI silencing. Moreover, an elevated IFNβ signature has been associated with more lymphocyte infiltration, better response to therapies, and better prognosis in TNBC (Cheon et al., 2023). An induced IFN response and elevated IFNβ expression were observed with CBP/P300 BRD inhibition *in vivo*. Both the IFN response and antigen presentation in tumor cells correlate with a better CTL response and high levels of tumor-infiltrated lymphocytes that have been associated with better prognosis and treatment response in TNBC (Huertas-Caro et al., 2023, Li and Cao, 2023). CBP/P300 BRD inhibitors were reported previously to reduce differentiation and an immunosuppressive phenotype of regulatory T cells in follicular lymphoma (Ghosh et al., 2016, Castillo et al., 2019). However, there is limited information about their effects on CTLs. One study suggested that CBP/P300 BRD inhibition may lead to a more activated phenotype in CTLs, but did not report an increase in CTL frequency in the CT26 tumors, a colon cancer and ‘hot tumor’ model known to have high lymphocyte infiltration and respond to ICB (Sato et al., 2021). In contrast, neutrophil-enriched TNBC are usually considered as “cold tumors”, inferring minimal lymphocyte infiltration and complete resistance to ICB (Bonaventura et al., 2019, Kim et al., 2019). In this study, we report that CBP/P300 BRD inhibition increased infiltrated CTLs in a “cold tumor” model and demonstrated that CTLs played a significant role in tumor regression and persistent growth inhibition induced by the CBP/P300 BRD inhibitor. These results imply that CBP/P300 BRD inhibition has the potential to reprogram “cold tumors” to “hot tumors”.

Importantly, in this study, we also tested the effects of the CBP/P300 BRD inhibitor in combination with chemotherapy and ICB. Docetaxel was selected because it was shown previously to stimulate immune response in both TNBC mouse models and patients (Vennin et al., 2023, Tsavaris et al., 2002). Taxanes and anti-PD-1 are a part of the standard-of-care regimen for early TNBC patients. In the current study, 2208L tumors exhibited a minimal response to docetaxel and anti-PD-1, but the addition of the CBP/P300 BRD inhibitor led to durable inhibition of tumor growth. Moreover, after 27 days of the treatment, we discontinued the CBP/P300 BRD inhibitor and docetaxel while continuing only anti-PD-1. The tumors in the combination treatment group displayed reduced tumor regrowth as compared to the single-agent treated group. In addition, TANs returned to the tumors treated with the single agent, but reduced TANs were still observed in the end-stage tumors from the combination treatment group. These findings suggest that the combination treatment may induce a more durable immune response. CTLs activated by anti-PD-1 treatment were previously reported to be capable of inducing apoptosis and inhibiting the activity of MDSCs (Chen et al., 2021). Thus, we speculate that the combination treatment might lead to a stronger and more persistent T-cell activation. These results suggest that adding a CBP/P300 BRD inhibitor to the standard-of-care therapies may help improve the response of neutrophil-enriched TNBC and result in a stronger immune response.

Lastly, we investigated the effects of CBP/P300 BRD inhibition on the treatment of established lung metastasis since metastasis is the cause of mortality for the majority of TNBC patients (Harbeck et al., 2019). In primary tumors, CBP/P300 BRD inhibition was found to reduce the RNA expression of multiple genes associated with metastasis. These findings implied that CBP/P300 BRD might play a role in metastasis. The 2208L model readily metastasizes to the lung, but 2208L tumors always exhibit aggressive local dissemination and invasion, and therefore we could never fully resect the primary tumors to study the treatment response of spontaneous metastases. Accordingly, in this study, we used an experimental metastasis model to investigate the effects of CBP/P300 BRD inhibition on established lung metastases.

CBP/P300 BRD inhibition significantly reduced the metastatic outgrowth of the 2208L model in the lung. Previous studies reported that neutrophils can support metastatic progression by inducing metastatic tumor cell proliferation and promoting angiogenesis (Leach et al., 2019).

CBP/P300 BRD inhibition was found to decrease infiltrated neutrophils in 2208L lung metastases. Thus, CBP/P300 BRD inhibitors may also be effective in treating metastases of neutrophil-enriched TNBC. This effect is also associated with reducing infiltrated neutrophils.

We are aware that our study has several limitations. All the models used in this study were transplantable syngeneic mouse models, and therefore surgeries were required to implant the tumors. These surgeries might trigger local inflammation and thus might transiently affect the TIME or the immune system systemically. Moreover, these models have relatively short latencies and rapid tumor growth, making it hard to assess treatment effects during earlier stages of tumor progression and T-cell responses and exhaustion which may require long-term treatment studies. Ideally, spontaneous autochthonous models could be used, but their long latency and variability preclude their use for treatment studies that require large and matched cohorts of control and treatment groups. Differences between the mouse and human immune systems might also affect the TIME. Since the response to CBP/P300 BRD inhibition is T cell-dependent, currently available immunocompromised PDX models are not suitable for these studies. To ensure that the responses observed were not dependent on the mouse strain, we selected models derived from both BALB/c and C57BL/6 backgrounds. Another limitation of the present study is that scRNA-seq does not infer changes in chromatin accessibility induced by CBP/P300 BRD inhibition. While single-cell ATAC sequencing coupled with scRNA-seq will provide more information such as changes in motif accessibility, due to the possibility of extracellular trap formation, neutrophils are routinely removed during sample preparation for single-cell ATAC sequencing.

In summary, this study suggests that CBP/P300 BRD inhibitors might provide therapeutic benefits to neutrophil-enriched TNBC by inhibiting immunosuppression and stimulating an antitumor immune response. With IACS-70654, a novel CBP/P300 BRD inhibitor, we determined that CBP/P300 BRD inhibition reduced the growth of neutrophil-enriched TNBC models. CBP/P300 BRD inhibition reduced TANs in the neutrophil-enriched model by reprogramming abnormal proliferation and differentiation of neutrophil progenitors in the bone marrow. CBP/P300 BRD inhibition also promoted IFN response, induced MHCI expression in tumor cells, and increased infiltrated CTLs. These results also indicated that the CBP/P300 BRD inhibitor may improve the response of neutrophil-enriched TNBC to chemotherapy and ICB, and the combination treatment might elicit a more durable immune stimulation. While these preclinical studies will need to be validated in patients, they provide a rationale for future clinical trials in neutrophil-enriched cancers.

## Methods

### IACS-70654 pharmacological characterization

Specific binding of the CBP or BRD4 bromodomain to the acetylated peptide derived from the H4 histone substrate (tetra acetylated H4 (1-21) Ac-K5/8/12/16) was measured in the absence or presence of inhibitors. The GST-tagged bromodomains of CBP (1081-1197) and BRD4 (49-170) were obtained from BPS Bioscience and binding to the biotinylated H4 (1-21) Ac-K5/8/12/16 (AnaSpec. 64989) was assessed via AlphaScreen technology (Perkin Elmer). For CBP AlphaScreen assay, 5 nM GST-CBP (1081-1197) and 20 nM biotin-H4 (1-21) Ac-K5/8/12/16 (AnaSpec. 64989) were incubated with varying concentrations of CBP inhibitors in 15 µl of buffer containing 50 mM HEPES 7.5, 100 nM NaCl, 1 mM TCEP, and 0.003% Tween-20. After 30 minutes of incubation at room temperature, 15 µl of detection buffer (BPS Bio. 33006) containing 7 mg/ml of Glutathione AlphaLisa acceptor beads (Perkin Elmer AL109) and 14 µg/ml of Streptavidin donor beads (Perkin Elmer 676002) was then added to the previous mixture. The reaction was incubated for an additional 2 hrs at room temperature, and the AlphaScreen signal was quantified using the Envision Multilabel plate reader. As a negative control, GST-CBP (1081-1197) was incubated with the non-acetylated biotin-H4 (1-21) peptide (AnaSpec. 62555) and in the presence of 0.25% of final DMSO concentration. For the BRD4 AlphaScreen assay, the binding of 2.5 nM of BRD4 (49-170) to 10 nM biotin-H4 (1-21) Ac-K5/8/12/16 (AnaSpec. 64989) was assessed following the same procedure described for the CBP assay. The standard dose response curves were fitted by Genedata Screener software using the variable-slope model. Only Signal and Dose in the equation were treated as known values. Screening of IACS-70654 against 32 bromodomain proteins was performed by BromoMAX (DiscoverX). To measure Kds, BromoKdELECT was used (DiscoverX).

### Animal Studies

The animal experiments were performed at Baylor College of Medicine following a protocol (AN-504) approved by the Institutional Animal Care and Use Committee. All female WT BALB/c mice were purchased from Inotiv, and all female WT C57BL/6J mice were purchased from The Jackson Laboratory. All animals were housed in the TMF mouse facility at Baylor College of Medicine with a 12-hr day or night cycle in climate-controlled conditions. All ethical regulations were complied with for all animal studies. The ethical point for primary mammary tumor study is tumor size greater than 1500 mm^3^. The ethical point for lung metastasis study is a more than 20% decrease in body weight or a significant decrease in body condition.

### Mammary tumor transplant

All *Trp53*-null tumor models were kept frozen in FBS + 10% DMSO as an established tumor bank in liquid nitrogen at the Rosen Lab. The generation of the *Trp53*-null tumor models has been described in previous publications (Jerry et al., 2000, Zhao et al., 2023). Before transplantation, frozen tumor pieces were thawed, washed with PBS, and cut into 1-mm chunks. Tumor transplantation was performed with 7-8 wks old WT BALB/c mice. One tumor chunk was implanted into the fourth mammary fat pad of each mouse. PyMT-N is a stable subtype derived from MMTV-PyMT tumors as described by our previous publication (Kim et al., 2019). PyMT-N tumors were kept frozen in FBS + 10% DMSO in liquid nitrogen at the Zhang Lab. The frozen PyMT-N tumor pieces were first transplanted to the fourth mammary fat pad of a 7–8-wk-old WT C57BL/6J mouse. When the tumor reached around 1 cm in diameter, it was harvested to obtain a freshly dissociated single-cell suspension. Tumors were chopped to fine paste using a McIlwain tissue chopper. Then tumors were digested with 1 mg/ml Collagenase Type I (Sigma-Aldrich, 11088793001) and 1 µg/ml DNase (Sigma-Aldrich, 11284932001) in DMEM/F12 media with no additives (GenDEPOT, CM020-050) for 90-120 minutes at 37°C on a shaker. The digested tumors were centrifuged at 1500 rpm, and the pellets were resuspended in PBS (GenDEPOT, CA008-050). To enrich tumor cells, the pellets underwent three 7-sec centrifugations at 1500 rpm, and the supernatants were discarded. Then the pellets were trypsinized, counted, filtered through a 40 µm cell strainer (VWR 352340), and resuspended in PBS. 250,000 PyMT-N tumor cells were injected into the mammary fat pad of each mouse. For all studies, tumor growth was monitored by measuring the tumor volume using a digital caliper. All treatment studies started when the average tumor volume was more than 80 mm^3^. The mice were randomized into treatment groups by weight and tumor volume using RandoMice (van Eenige et al., 2020). Then weight and tumor volume measurements were performed 3 times a week during the treatment studies. The animal studies were not blinded, but for some studies, tumor volume was measured by two independent persons.

### Experimental lung metastasis model using tail vein injection

To prepare for the tail vein injection, a 2208L tumor chunk was first implanted into the fourth mammary fat pad of a 7–8-wk-old WT BALB/c mouse. When the tumor reached around 1 cm in diameter, the tumor was harvested and processed into a single-cell suspension as described above with PyMT-N tumors. Lung metastases were generated via tail vein injection of 100,000 cells into each 6-8 wks-old female BALB/c mouse using a 27-gauge syringe. Because the tumor cells were not labeled, two mice were euthanized to collect the lung to ensure a successful generation of lung metastases before starting the treatment studies.

### Treatment

IACS-70654 was suspended in sterile 0.5% methylcellulose (Sigma-Aldrich, M0430) and administered orally at the dosage of 3.75 mg/kg on a 3-on/2-off schedule. The control group was given the same volume of 0.5% methylcellulose. For labeling proliferative cells, mice were injected i.p. with 60 mg/kg BrdU (Sigma-Aldrich, B-5002) 2 hrs before euthanasia. Docetaxel (DTX) (LC Laboratories, D-1000) was dissolved in Tween 80 first and then diluted 1:4 with 16.25% ethanol. DTX was administered i.p. at 10 mg/kg weekly. RecombiMAb anti-mouse PD-1 (BioXCell, CP151) and RecombiMAb mouse IgG2a isotype (BioXCell, CP150) were administered i.p. at 200 µg per mouse every 3 days. InVivoMAb anti-mouse CD8α (BioXCell, BE0061) and InVivoMAb rat IgG2b isotype control (BioXCell, BE0090) were administered i.p. at 250 µg and 200 µg per mouse respectively every four days. All *in vivo* antibodies were diluted with InVivoPure pH 7.0 Dilution Buffer (BioXCell, IP0070). Treatment studies of IACS-70654 as a single agent were repeated at least twice (performed 3 times in total). Studies of combination treatments were repeated once (performed twice in total). The mice were treated in a random order during each treatment. Mice were monitored for signs of toxicity at least 3 times per week.

## Flow cytometry

### Tumor-infiltrated immune cells

Tumors were processed and digested with collagenase as previously described. To enrich immune cells, the supernatants were collected after the three 7-sec centrifugations. The supernatants were spun down, resuspended in RBC lysis buffer (BioLegend, 420301), and incubated on ice for 2 minutes. The enriched cells were spun down, resuspended in DMEM/F12 + 2% FBS, and filtered through a 40 µm cell strainer. Live cells were counted using a Bio-Rad TC20 Cell Counter with 0.4% Trypan Blue stain (Gibco, 15250-061). Cells were spun down and resuspended to 10 million cells/ml in FACS buffer (PBS + 2% FBS). 1.0 x 10^6^ cells from each sample were loaded onto a 96-well non-treated tissue culture plate (CELLTREAT, 229590).

In a 96-well non-treated plate, 1.0 x 10^6^ cells were stained with Live/Dead Fixable Yellow (ThermoFisher Scientific L34968, 1:800). The cells were washed with FACS buffer and blocked with anti-mouse CD16/32 antibody (BioLegend, 101320). Then the cells were stained with antibodies for cell surface markers from each panel. Panel 1 includes MHCII BV711 (BioLegend, 107643, 1:500), CD11b PerCP-Cy5.5 (BioLegend, 101227, 1:400), Ly6C PE-Cy7 (BioLegend, 128017, 1:300), F4/80 APC (BioLegend, 123115, 1:150), CD45 Alexa Fluor 700 (BioLegend, 103128, 1:250) and Ly6G APC/Fire 750 (BioLegend, 127651, 1:200). Panel 2 includes CD45 Pacific Blue (BioLegend, 103126, 1:300), CD3ɛ PerCP-Cy 5.5 (BioLegend, 100218, 1:100), CD45R (B220) PE-Cy7 (BioLegend, 103221, 1:200), CD8α Alexa Fluor 700 (BioLegend, 100730, 1:200), CD4 APC/Fire 750 (BioLegend, 100460, 1:100) and CD279 (PD-1) BV711 (BioLegend, 135231, 1:150). For intracellular staining, the cells were fixed and permeabilized using the FOXP3/Transcription Factor Staining Buffer Set (ThermoFisher Scientific,00-5523-00). The cells were then blocked with 2% rat serum (Sigma-Aldrich, R9759) and 2% goat serum (Sigma-Aldrich, G9023) in permeabilization buffer from the buffer set. The cells were then stained with antibodies for intracellular markers from each panel. Panel 1 includes Arginase PE (Fisher Scientific, 12-3697-82, 1:200) and CD206 PE/Dazzle 594 (BioLegend, 141732, 1:150). Panel 2 includes Foxp3 PE (BioLegend, 126403, 1:150) and Granzyme B APC (BioLegend, 372203, 1:150). For detecting Acetyl-histone H3, the cells were first incubated with Acetyl-histone H3 (Lys27) antibody (Cell Signaling, 8173T, 1:200) with antibodies for other intracellular markers. Subsequently, the cells were stained with the anti-rabbit secondary antibody conjugated with Alexa Fluor 488 (Invitrogen, A-11008) or AlexaFluor 594 (Invitrogen, A-11012) at 1:500 dilution for 1 hr. After staining, the cells were washed with permeabilization buffer and resuspended in PBS for acquisition on the Attune NxT flow cytometer at the Baylor College of Medicine FACS and Cell Sorting core. The collected data was compensated and analyzed using FlowJo v10 software (FlowJo, RRID: SCR_008520).

### Blood and bone marrow myeloid cells

For PBMC, blood was collected retro-orbitally, and 30 µL of blood from each mouse was incubated with 3 ml of RBC lysis buffer for 30 minutes. For the bone marrow, the mice were injected i.p. with 60mg/kg BrdU 24 hrs before euthanasia. Then bone marrow cells were extracted from one femur bone of each mouse. The cells were incubated with 2 ml of RBC lysis buffer for 2 minutes and then filtered through a 40-µm cell strainer. Bone marrow cells were stained with Live/Dead Fixable Yellow at 1:800 dilution. To prepare for marker staining, all samples were washed with FACS buffer and blocked with anti-mouse CD16/32 antibody in a 96-well non-treated plate on ice for 15 minutes. The cells were then incubated with antibodies for including CD45 FITC (BioLegend, 103108, 1:500), CD11b PerCP-Cy5.5 (BioLegend, 101227, 1:500), F4/80 APC (BioLegend, 123115, 1:150), Ly-6C PE-Cy7 (BioLegend, 128017, 1:300), and Ly6G APC/Fire 750 (BioLegend, 127651, 1:200). PBMC were washed and stained with NucBlue Live ReadyProbes Reagent (ThermoFisher Scientific, R37605) in PBS for data acquisition on the flow cytometer. The bone marrow cells were fixed and permeabilized using the FOXP3/Transcription Factor Staining Buffer Set overnight. On the next day, the bone marrow cells were washed with permeabilization buffer and incubated in 0.3 mg/ml DNase I at 37 °C for 1 hr. Then the bone marrow cells were washed and stained with BrdU PE (BioLegend, 339812, 1:100) for 30 minutes. After washing, the bone marrow cells were resuspended in PBS for data acquisition on the flow cytometer. The collected data was compensated and analyzed using FlowJo v10 software.

## Histology

Primary tumor and lung tissues were fixed in 4% Paraformaldehyde overnight and then stored in 70% ethanol. The fixed tissues were embedded in paraffin and sectioned at 5-µm at the Breast Center Pathology Core at Baylor College of Medicine. Serial sectioning of lung metastases was performed by collecting 4-6 5-µm sections every 150 µm.

### Immunohistochemical (IHC) staining

The tissue sections were first deparaffinized and rehydrated using xylene and 100%, 95%, 80%, and 75% ethanol. Antigen retrieval was performed by immersing tissue sections in Tris-EDTA antigen retrieval buffer (10 mM Tris base, 1 mM EDTA, 0.05% Tween-20, pH 9.0) with heating on a steamer for 30 minutes. Then the sections were washed, and endogenous peroxidases were blocked by immersing tissue sections in 3% hydrogen peroxide (Fisher Scientific, H323-500, diluted 1:10 by PBS) for 10 minutes. The sections were washed and incubated for 1 hr in the IHC blocking buffer containing 3% BSA (Sigma-Aldrich, A7906) and 5% goat serum (Sigma-Aldrich, G9023) in PBS. After blocking, tissue sections were incubated with primary antibodies at 4°C overnight. The primary antibodies used include S100A8 (R&D Systems, MAB3059, 1:5000), F4/80 (Cell Signal Technology, 70076S, 1:500), BrdU (Abcam, ab6326, 1:1000) and CD8α (Cell Signal Technology, 98941S, 1:500). The sections were washed and then incubated in biotin-conjugated anti-rat (Vector Laboratories, PI-9400-1) or anti-rabbit (Vector Laboratories, PI-1000-1) secondary antibodies at 1:1000 dilution for 1 hr at room temperature. Next, sections were incubated with VECTASTAIN Elite ABC HRP Reagent (Vector Laboratories, PK7100) for 30 min and treated with ImmPACT DAB peroxidase substrate (Vector Laboratories, sk-4105) until optimal signals were observed. The slides were counterstained with Harris Hematoxylin with Glacial Acetic Acid (Poly Scientific, S212A). Then the slides were dehydrated by incubation at 60 °C for 1 hr and mounted in Poly-Mount Xylene (Poly Scientific, 24176–120).

### H&E staining

After deparaffinization and rehydration, the slides were stained with Harris Hematoxylin with Glacial Acetic Acid and Eosin Phloxine Alcoholic Working Solution (Poly Scientific, S176). The slides were then dehydrated using 95% ethanol, 100% ethanol, and xylene. Then the slides were mounted in Poly-Mount Xylene.

### Imaging and quantification

Images of all IHC or H&E-stained slides were taken using the Olympus BX40 light microscope and MPX-5C pro low light camera at 20X magnification or with Aperio ImageScope (Leica Biosystems). Quantification was performed by either Fiji (Schindelin et al., 2012) using 3-5 microscope images per section.

## Single-cell RNA sequencing

### Single-cell suspension preparation for single-cell RNA sequencing

For the tumor single-cell RNA sequencing (scRNA-seq), tumors were processed as described previously and dissociated with Collagenase Type I. A shortened Collagenase (1 hr) was performed to maximize cell viability. The tumor digest was centrifuged at 1500 rpm, and then the pellet was incubated in 2 ml RBC lysis buffer for 2 minutes at room temperature. The cells were filtered through a 40-µm cell strainer and resuspended in PBS. Three mice were used for each treatment arm, and the cells from the three mice were pooled. Dissociated cells were stained with Ghost Dye UV 450 (Tonbo Biosciences, 13-0868) for 10 minutes on ice. Next, dissociated cells were centrifuged, resuspended in FACS buffer, and FACS-sorted to select viable cells (Ghost Dye UV 450-) for analysis.

For the bone marrow scRNA-seq, bone marrow cells were harvested from the tibia and femur bones using PBS (2% FBS) and then passed through a 70 µm strainer. Following centrifugation at 600g for 5 min, the cells were resuspended in red blood cell lysis buffer (Tonbo Biosciences, SKU TNB-4300-L100) for 10 min at room temperature. To isolate bone marrow CD45+Ter119-cells, total bone marrow cells were stained with anti-mouse CD45-APC (Tonbo Biosciences, 20-0451-U100), anti-mouse Ter119-PE (BioLegend,116207) and DAPI (Invitrogen, R37606) at 4°C for 15 min, followed by FACS-sorting of bone marrow DAPI-CD45+Ter119-cells. To isolate bone marrow HSPCs, total bone marrow cells were processed as above steps, then stained with anti-mouse biotinylated-lineage antibodies (CD11b/Gr-1/B220/Ter119/CD3e) (BD Bioscience, 559971), followed by staining of Streptavidin-APC (Tonbo Biosciences, 20-4317), CD45-VF450 (Tonbo Biosciences, 75-0451), c-Kit-PE/Cy7 (Tonbo Biosciences, 60-1172) and DAPI. The DAPI-CD45+Lin-c-Kit+ (enriched HSPCs) were then FACS-sorted for analysis.

The single-cell suspensions from tumor or bone marrow were then tagged with cellplex barcoding oligo (10x Genomics, 1000261), and immediately delivered to the Single Cell Genomics Core at Baylor College of Medicine for scRNA-Seq library preparation.

### Library preparation and single-cell RNA sequencing

The single-cell gene expression Library was prepared according to the Chromium Single Cell Gene Expression 3’v3.1 kit along with the feature barcoding kit (10x Genomics, 1000262).

Briefly, single cells, reverse transcription reagents, Gel Beads containing barcoded oligonucleotides, and oil were loaded on a Chromium controller (10x Genomics) to generate single-cell GEMS (Gel Beads-In-Emulsions) where full-length cDNA was synthesized and barcoded for each single cell. Subsequently, the GEMS are broken and cDNA from each single cell is pooled. Following cleanup using Dynabeads MyOne Silane Beads, cDNA is amplified by PCR. The amplified product is fragmented to optimal size before end-repair, A-tailing, and adaptor ligation. The final GEX and Cellplex library were generated by amplification. After passing the quality control, the next-generation sequencing of libraries was performed on NovaSeq 6000 (Illumina).

### Pre-processing of scRNA-seq datasets

Upon receiving the raw sequencing data, we prepared the cellranger multi (v7.2.0) pipeline to conduct alignment, read counts, and sample demultiplexing. For the tumor dataset, we set the barcode assignment confidence as 0.8 to include more cells for downstream analysis. Downstream analyses of single-cell RNA-seq were performed using the Seurat (v4.4.0) package in R (R version 4.3.1). For quality control, we kept cells with more than 200 and less than 6000 read counts. Cells with more than 1000 UMI counts and more than 10% mitochondrial ratio were also removed. Datasets were downsampled to the lowest cell number in the group (tumor, bone marrow CD45+Ter119-or HSPC) and normalized. The top 2000 most variable genes were selected for each sample. Anchors among all samples were generated with the FindIntegrationAnchors function in Seurat, and then the samples were integrated with the anchors using the IntegrateData function in Seurat.

### Clustering and cluster annotation

The integrated data was used to perform principal component analysis (PCA). Then the first 30 principal components identified by PCA were used for Uniform Manifold Approximation and Projection (UMAP) analysis. For the tumor sample, cells were clustered at a resolution of 0.5. The clusters were annotated using SingleR Immgen and then verified manually and by the FindMarkers function. The macrophage and monocyte populations are isolated and re-clustered at a resolution of 0.4. The CD45+ bone marrow cells were clustered at a resolution of 0.5. The clusters were first annotated using SingleR Immgen, and only the myeloid clusters (Neutrophils, Monocytes, and dendritic cell progenitors) were kept for analysis. Then the neutrophils were isolated and re-clustered at the resolution of 0.7. Clusters of neutrophils were annotated manually based on published markers (Grieshaber-Bouyer et al., 2021, Carnevale et al., 2023, Qu et al., 2023) and markers identified using the FindMarkers function. Some clusters were combined if needed. For HSPCs, clustering was performed at the resolution of 1.2, and non-HSPC clusters were excluded based on markers published in the previous publication (Hao et al., 2023). The HSPCs were re-clustered at the resolution of 1 and re-annotated using published markers (Hao et al., 2023). All pseudotime plots were generated using Monocle 3, and the clustering was performed at the resolution of 1e-3. The cell cycle scoring was performed using the CellCycleScoring function in Seurat, and the cell cycle gene signatures were converted from human genes to mouse genes by using the gorth function in the gprofiler2 package.

### Differentially Expressed Genes

Differentially expressed genes were identified with FindMarkers function from Seurat using the MAST test between two groups. The volcano plot was generated using the EnhancedVolcano package. The clusterProfiler package was used for GO analyses of the tumor cell population with an adjusted p-value cutoff smaller than 0.01 and a log2 fold change cutoff greater than 0.5. Enriched pathways were identified with a p-value cutoff of 0.05, and the minimum gene count was 5. The enriched pathways were also simplified to minimize repetitive pathways.

## Immunoblotting

Tumors were snap-frozen and later homogenized in tissue lysis buffer (62.5 mM Tris-HCl pH 6.8, 2% SDS) containing cOmplete, EDTA-free Protease Inhibitor Cocktail (Sigma-Aldrich, 11873580001) with zirconium beads (Benchmark Scientific, D1132-30,) in BeadBlaster 24 Microtube Homogenizer. The homogenate was heated at 98°C for 8 minutes, and the protein concentrations were measured with the BCA Protein Assay Kit (ThermoFisher, 23227). The protein extracts were loaded on an SDS-PAGE system and then transferred to polyvinylidene difluoride membranes (Millipore, IPVH00010). The primary antibodies used include FGFR1 (Cell Signaling Technology, 9740, 1:1000) and β-Actin (Cell Signaling Technology, 3700, 1:5000).

## Cytokine Profiling

Tumor homogenates were prepared from tumors snap-frozen at the time of harvest. Tumors were homogenized with T-PER Tissue Protein Extraction Reagent (ThermoScientific, 78510) containing cOmplete, EDTA-free Protease Inhibitor Cocktail with zirconium beads in BeadBlaster 24 Microtube Homogenizer. Protein concentrations of the homogenates were measured with BCA Protein Assay Kit, and all tumor homogenates were diluted to 1.8 µg/µL. Plasma was collected by centrifuging blood samples twice at 1000 x g for 10 minutes. All plasma samples were diluted two-fold with PBS. All tumor homogenate and plasma samples were assayed using Mouse Cytokine/Chemokine 44-Plex Discovery Assay Array (MD44) at Eve Technologies Corp.

## Statistical Analysis

Sample sizes for all animal studies were determined based on preliminary studies. For comparison between 2 groups, unpaired, two-tailed Student’s *t* tests were used. For comparison among 3 or more groups and pairwise comparisons, ordinary one-way ANOVA and Tukey’s multiple comparisons were used. Two-way ANOVA and Sidak’s multiple comparisons were used for analyzing tumor volume changes, log fold changes of tumor volume or body weight changes over time. For Kaplan-Meier curves, the log-rank test was performed, and for multiple comparisons of survival analyses, the Bonferroni method was used to adjust the P-value. A P-value less than 0.05 was considered significant. All statistical analyses mentioned above were performed with GraphPad Prism 9. For all single-cell RNA-seq analyses, statistical analyses were performed in R, and an adjusted P-value less than 0.01 was considered significant.

## Online supplemental material

Tables S1 and 2 show the potency and selectivity of IACS-70654. Supplemental Figure S1 shows the structure of IACS-70654 and supplemental data of IACS-70654 7-day treatment. Figure S2 shows supplemental data on changes in tumor-associated myeloid cells. Figure S3 shows supplemental scRNA-seq analyses of bone marrow neutrophils. Figure S4 shows supplemental scRNA-seq analyses of bone marrow HSPCs. Figure S5 shows supplemental scRNA-seq analyses of 2208L tumors. Figure S6 includes supplemental data of IACS-70654 combination studies with DTX, anti-PD-1, and anti-CD8. Figure S7 includes supplemental analyses of metastasis-related genes and data of 2208L experimental lung metastases.

## Data availability

Raw and processed data of all single-cell RNA sequencing experiments including the metadata have been submitted to the GEO database and are publicly available (GSE264627). All other data or experiment information from this study are available in the article or the online supplemental material and from the corresponding author upon request.

## Acknowledgments

We thank Yang Gao at Baylor College of Medicine for providing the PyMT-N tumor model. We thank the Breast Center Pathology Core at Baylor College of Medicine for technical support. J.M.R. is supported by NCI CA148761. N.Z. is supported by CPRIT RP220468. D.A.P. is supported by American Cancer Society PF-22-163-01-MM. A.J.S. is supported by Komen SAC232150. X.H.-F.Z. is supported by the US Department of Defense DAMDW81XWH-16-1-0073 (Era of Hope Scholarship), NCI (CA183878, CA251950, CA221946, CA227904, and CA253533), DAMD W81XWH-20-1-0375, Breast Cancer Research Foundation, and McNair Medical Institute. Flow cytometry analysis and cell sorting were supported by the Cytometry and Cell Sorting Core at Baylor College of Medicine with funding from the CPRIT Core Facility Support Award (CPRIT-RP180672), the NIH (CA125123 and RR024574) and the assistance of Joel M. Sederstrom. The scRNA-seq experiments were performed at the Single Cell Genomics Core at BCM partially supported by NIH S10OD025240, and CPRIT RP200504. IACS-70654 is covered by Patent US10899769, and K.L., M.J.S., and P.J. are inventors of the patent. The graphic abstract is created with BioRender.com. The authors declare no competing financial interests.

