## Supplemental Tables and Figures for "CBP/P300 BRD Inhibition Reduces Neutrophil Accumulation and Activates Antitumor Immunity in TNBC"

| | IACS-70654 (nM) |
|---|---|
| **CBP $IC_{50}$** | 5.5 |
| **BRD4 $IC_{50}$** | 544 |

**Supplementary Table S1.** Specific binding of the CBP or BRD4 bromodomain by IACS-70654.

| Target | Kd (nM) | Selectivity |
|---|---|---|
| **CBP** | 0.096 | 1x |
| **P300** | 0.15 | 1.6x |
| **BRD2(1)**<br>**BRD2(2)** | 17<br>390 | 177x<br>4062x |
| **BRD3(1)**<br>**BRD3(2)** | 45<br>89 | 469x<br>927x |
| **BRD4(1)**<br>**BRD4(2)** | 24<br>210 | 250x<br>2188x |
| **BRDT(1)**<br>**BRDT(2)** | 7.5<br>170 | 78x<br>1771x |
| **WDR9(2)** | 16 | 167x |

**Supplementary Table S2.** Kd and selectivity of IACS-70654 against 32 bromodomain proteins.

### Figure S1

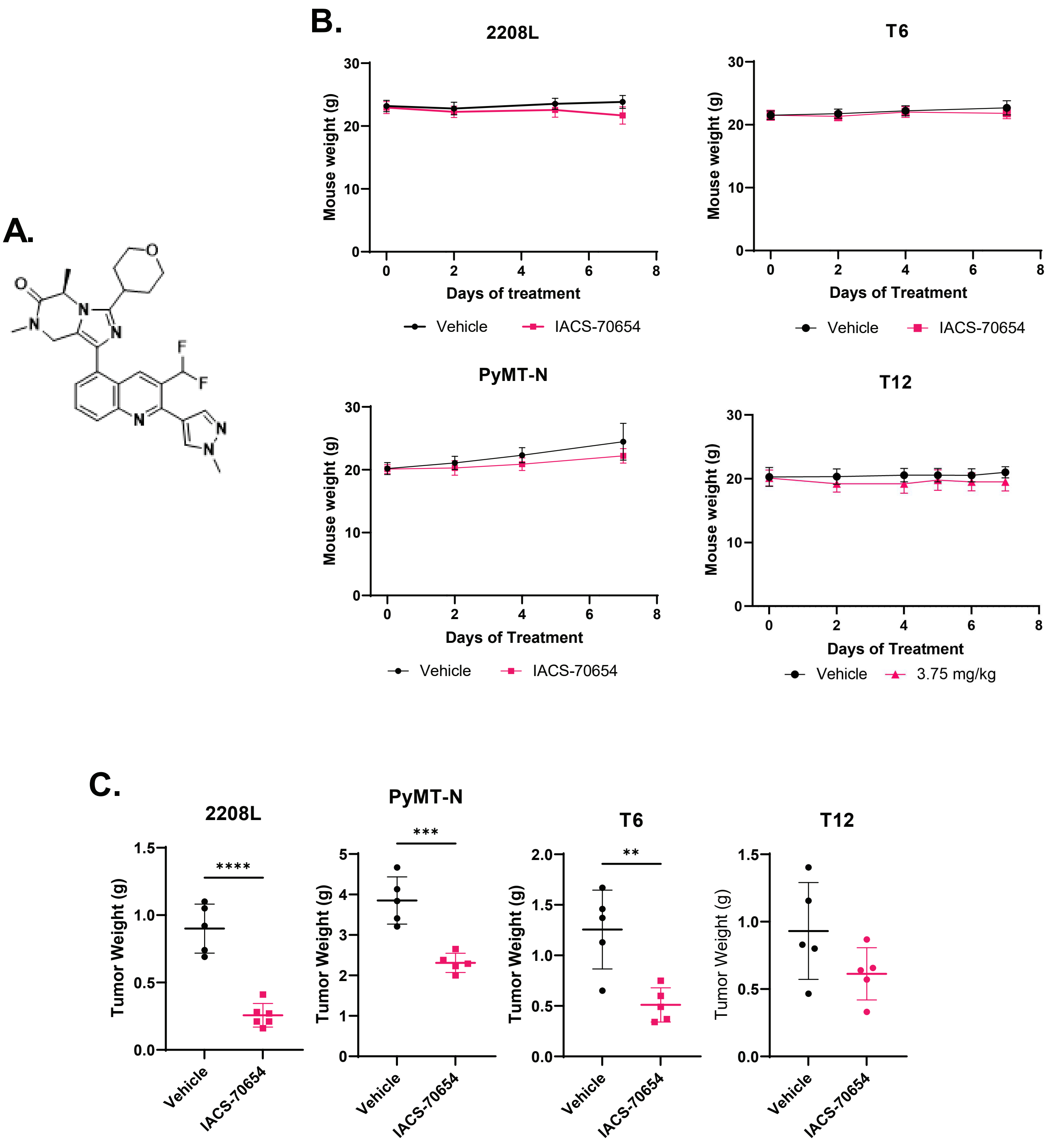

A.
O
N
N
N
F
F
N
N
N
O
B.
2208L
Mouse weight (g)
30
20
10
0
0
2
4
6
8
Days of treatment
Vehicle
IACS-70654
T6
Mouse weight (g)
Days of Treatment
Vehicle
IACS-70654
PyMT-N
Mouse weight (g)
Days of Treatment
Vehicle
IACS-70654
T12
Mouse weight (g)
Days of Treatment
Vehicle
3.75 mg/kg
C.
2208L
Tumor Weight (g)
1.5
1.0
0.5
0.0
****
Vehicle
IACS-70654
PyMT-N
Tumor Weight (g)
5
4
3
2
1
0
***
Vehicle
IACS-70654
T6
Tumor Weight (g)
2.0
1.5
1.0
0.5
0.0
**
Vehicle
IACS-70654
T12
Tumor Weight (g)
1.5
1.0
0.5
0.0
Vehicle
IACS-70654

### Figure S2

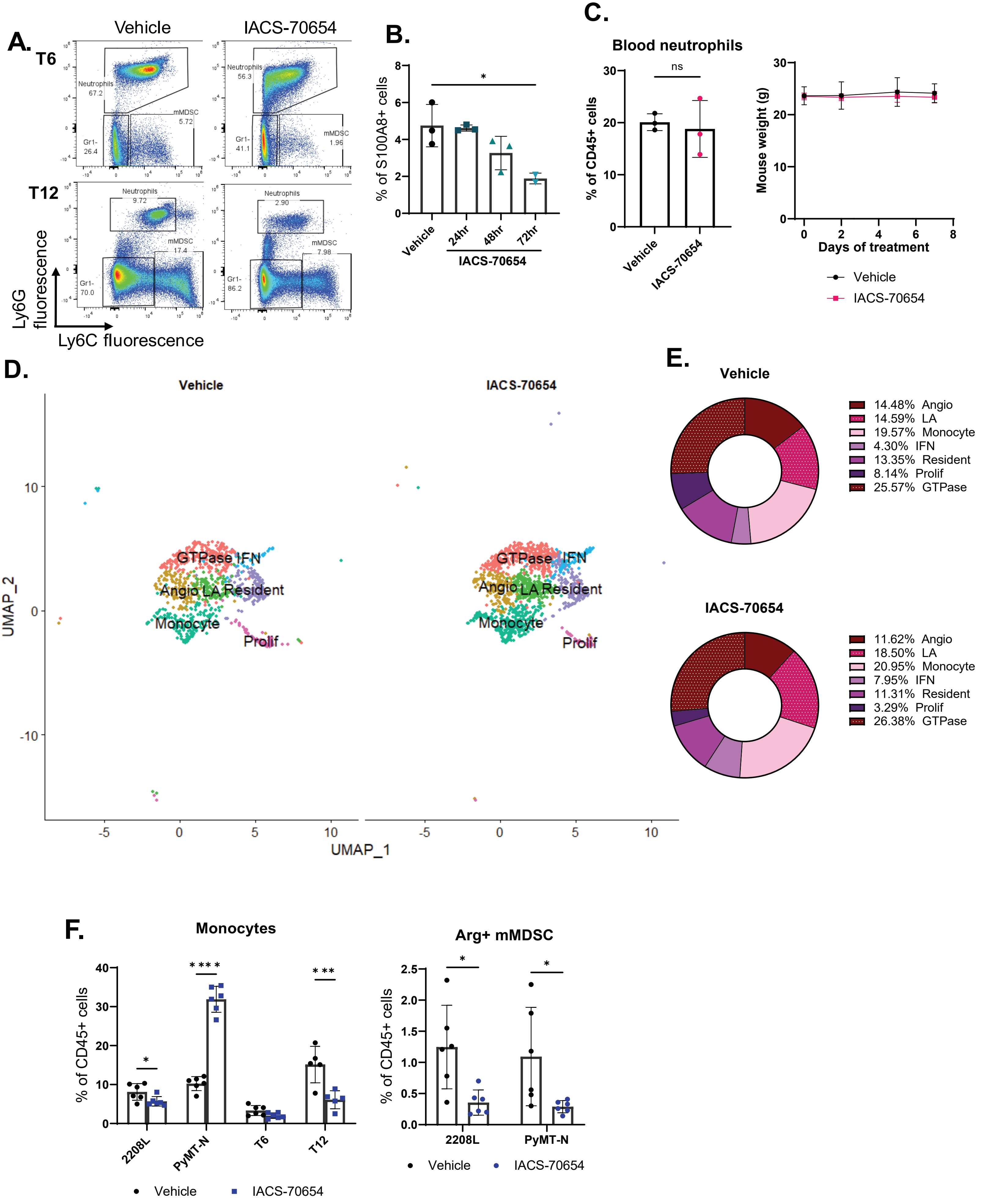

A.
Vehicle
IACS-70654
T6
T12
Neutrophils 67.2
mMDSC 5.72
Gr1- 26.4
Neutrophils 56.3
mMDSC 1.96
Gr1- 41.1
Neutrophils 9.72
mMDSC 17.4
Gr1- 70.0
Neutrophils 2.90
mMDSC 7.98
Gr1- 86.2
Ly6G fluorescence
Ly6C fluorescence
B.
% of S100A8+ cells
Vehicle
24hr
48hr
72hr
IACS-70654
C.
Blood neutrophils
ns
% of CD45+ cells
Vehicle
IACS-70654
Mouse weight (g)
Days of treatment
Vehicle
IACS-70654
D.
Vehicle
IACS-70654
GTPase
IFN
Angio
LA
Resident
Monocyte
Prolif
UMAP_2
UMAP_1
E.
Vehicle
14.48% Angio
14.59% LA
19.57% Monocyte
4.30% IFN
13.35% Resident
8.14% Prolif
25.57% GTPase
IACS-70654
11.62% Angio
18.50% LA
20.95% Monocyte
7.95% IFN
11.31% Resident
3.29% Prolif
26.38% GTPase
F.
Monocytes
% of CD45+ cells
2208L
PyMT-N
T6
T12
Vehicle
IACS-70654
Arg+ mMDSC
% of CD45+ cells
2208L
PyMT-N
Vehicle
IACS-70654

### Figure S3

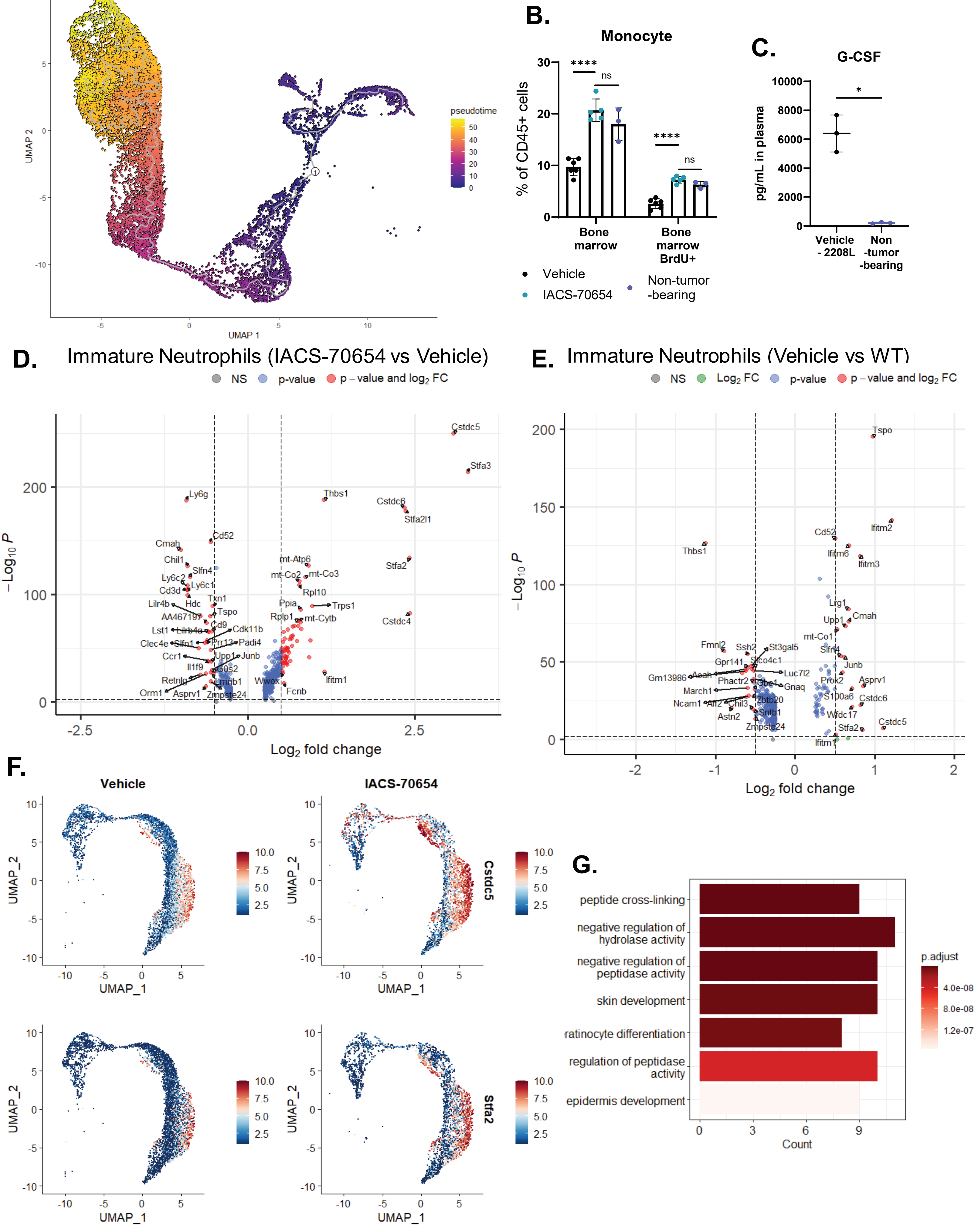

A.
pseudotime
UMAP 1
UMAP 2
B.
Monocyte
% of CD45+ cells
Bone marrow
Bone marrow BrdU+
Vehicle
IACS-70654
Non-tumor -bearing
C.
G-CSF
pg/mL in plasma
Vehicle - 2208L
Non -tumor -bearing
D.
Immature Neutrophils (IACS-70654 vs Vehicle)
NS
p-value
p – value and log2 FC
Log2 fold change
E.
Immature Neutrophils (Vehicle vs WT)
NS
Log2 FC
p-value
p – value and log2 FC
Log2 fold change
F.
Vehicle
IACS-70654
Cstdc5
Stfa2
UMAP_1
UMAP_2
G.
peptide cross-linking
negative regulation of hydrolase activity
negative regulation of peptidase activity
skin development
ratinocyte differentiation
regulation of peptidase activity
epidermis development
p.adjust
Count

### Figure S4

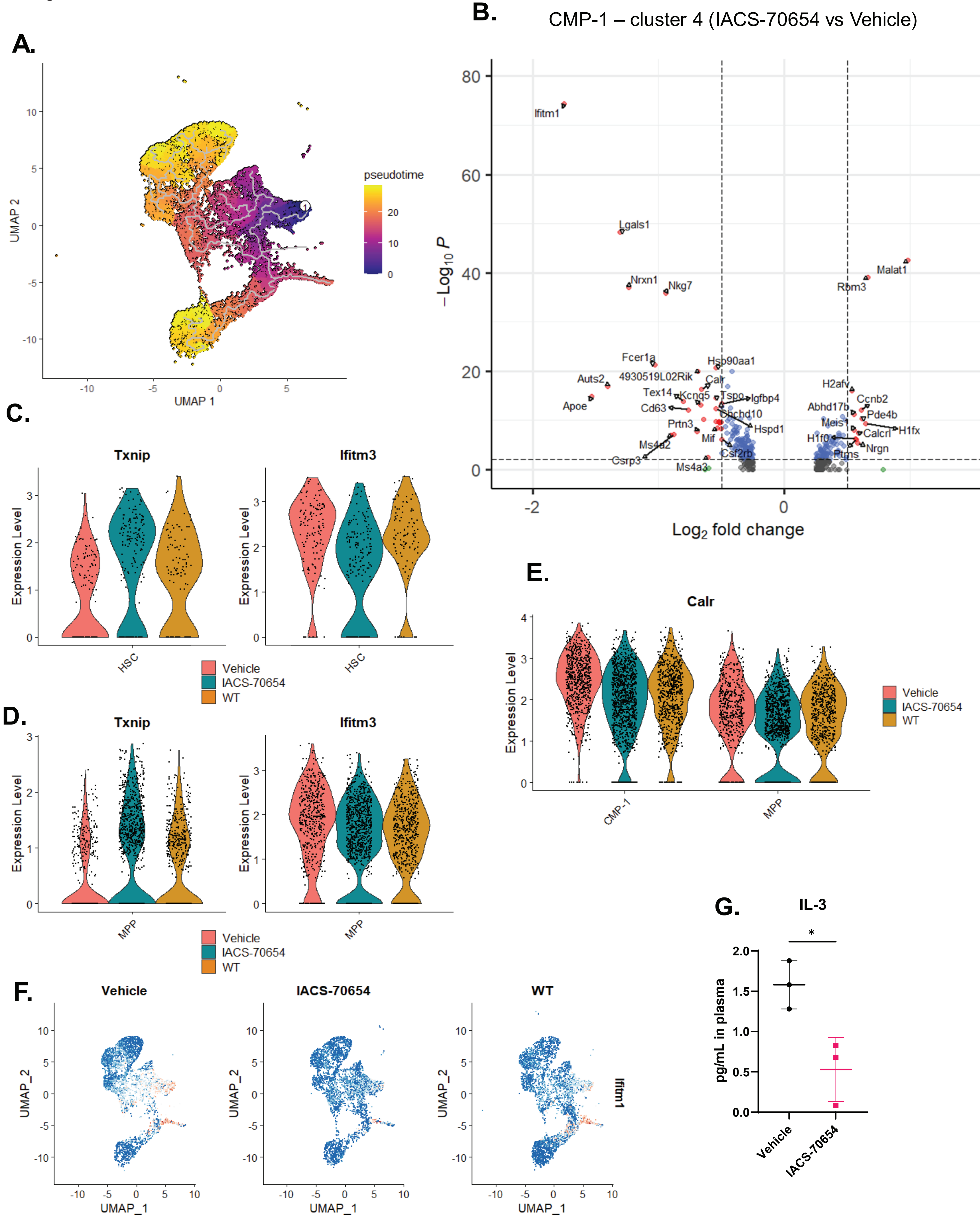
A.
pseudotime
UMAP 2
UMAP 1
B.
CMP-1 – cluster 4 (IACS-70654 vs Vehicle)
Ifitm1
Lgals1
Nrxn1
Nkg7
Malat1
Rbm3
Fcer1a
Hsp90aa1
Auts2
4930519L02Rik
Calr
Tex14
Kcnq5
Tspo
Igfbp4
Apoe
Cd63
Chchd10
Prtn3
Hspd1
Mif
Ms4a2
Csf2rb
Csrp3
Ms4a3
H2afv
Ccnb2
Abhd17b
Pde4b
Meis1
H1fx
H1f0
Calcrl
Nrgn
Ptms
$-Log_{10}$ P
$Log_2$ fold change
C.
Txnip
Ifitm3
Expression Level
HSC
Vehicle
IACS-70654
WT
D.
Txnip
Ifitm3
Expression Level
MPP
Vehicle
IACS-70654
WT
E.
Calr
Expression Level
CMP-1
MPP
Vehicle
IACS-70654
WT
F.
Vehicle
IACS-70654
WT
UMAP_2
UMAP_1
Ifitm1
G.
IL-3
pg/mL in plasma
*
Vehicle
IACS-70654

Figure S5

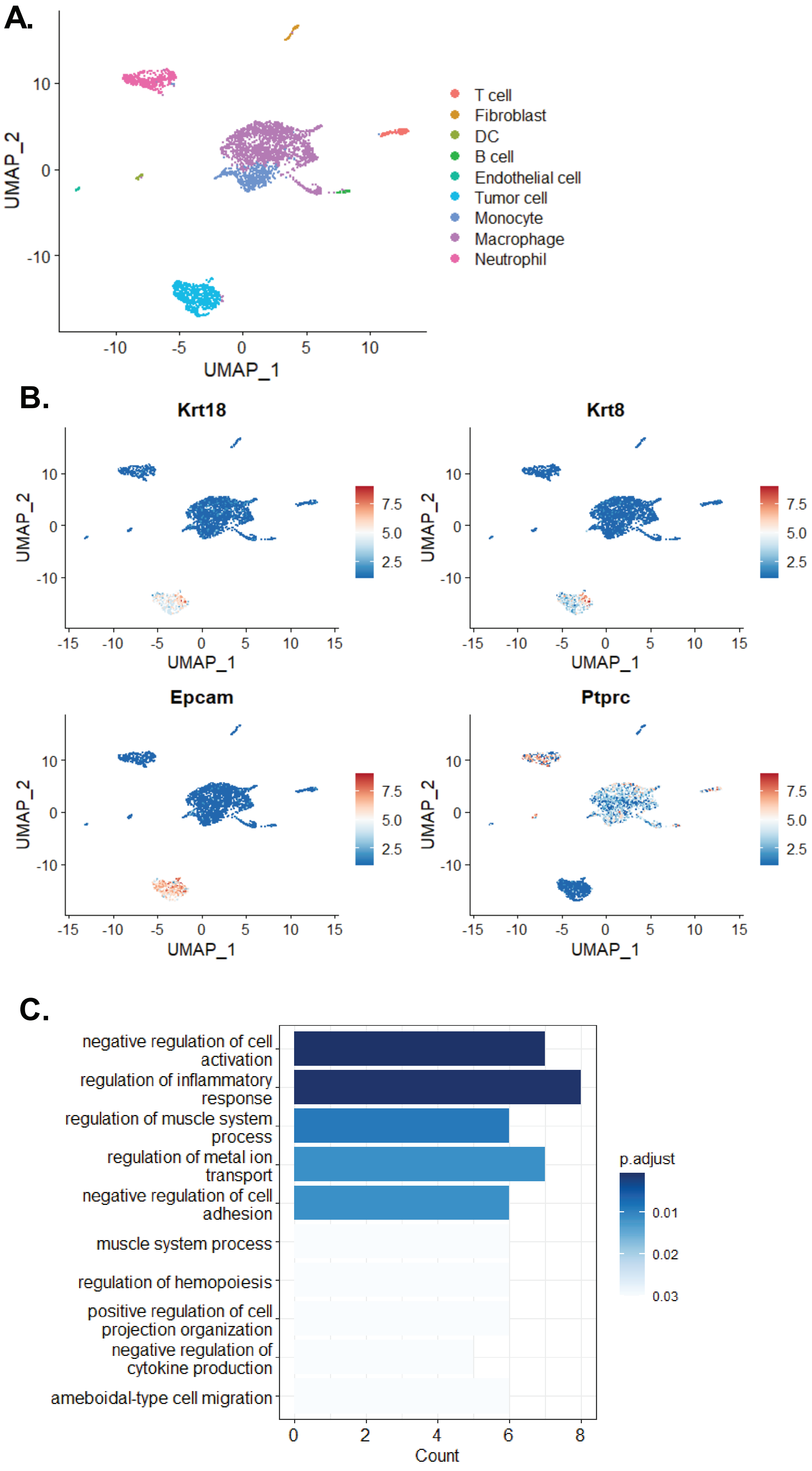
A.
UMAP_2
UMAP_1
T cell
Fibroblast
DC
B cell
Endothelial cell
Tumor cell
Monocyte
Macrophage
Neutrophil
B.
Krt18
Krt8
Epcam
Ptprc
C.
negative regulation of cell activation
regulation of inflammatory response
regulation of muscle system process
regulation of metal ion transport
negative regulation of cell adhesion
muscle system process
regulation of hemopoiesis
positive regulation of cell projection organization
negative regulation of cytokine production
ameboidal-type cell migration
Count
p.adjust

Figure S6

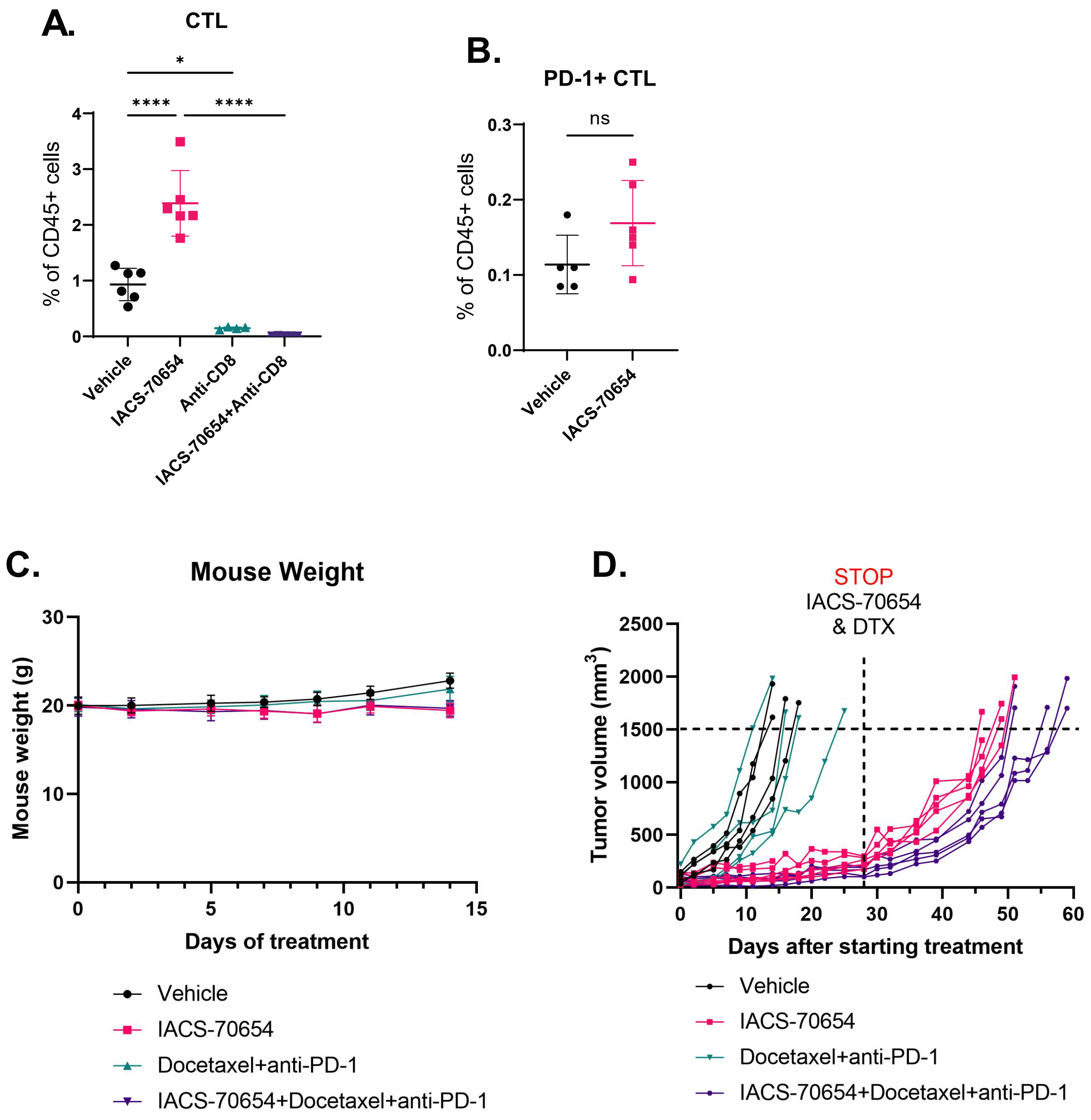
A.
CTL
*
****
****
% of CD45+ cells
0
1
2
3
4
Vehicle
IACS-70654
Anti-CD8
IACS-70654+Anti-CD8
B.
PD-1+ CTL
ns
% of CD45+ cells
0.0
0.1
0.2
0.3
Vehicle
IACS-70654
C.
Mouse Weight
Mouse weight (g)
0
10
20
30
0
5
10
15
Days of treatment
Vehicle
IACS-70654
Docetaxel+anti-PD-1
IACS-70654+Docetaxel+anti-PD-1
D.
STOP
IACS-70654
& DTX
Tumor volume (mm3)
0
500
1000
1500
2000
2500
0
10
20
30
40
50
60
Days after starting treatment
Vehicle
IACS-70654
Docetaxel+anti-PD-1
IACS-70654+Docetaxel+anti-PD-1

Figure S7

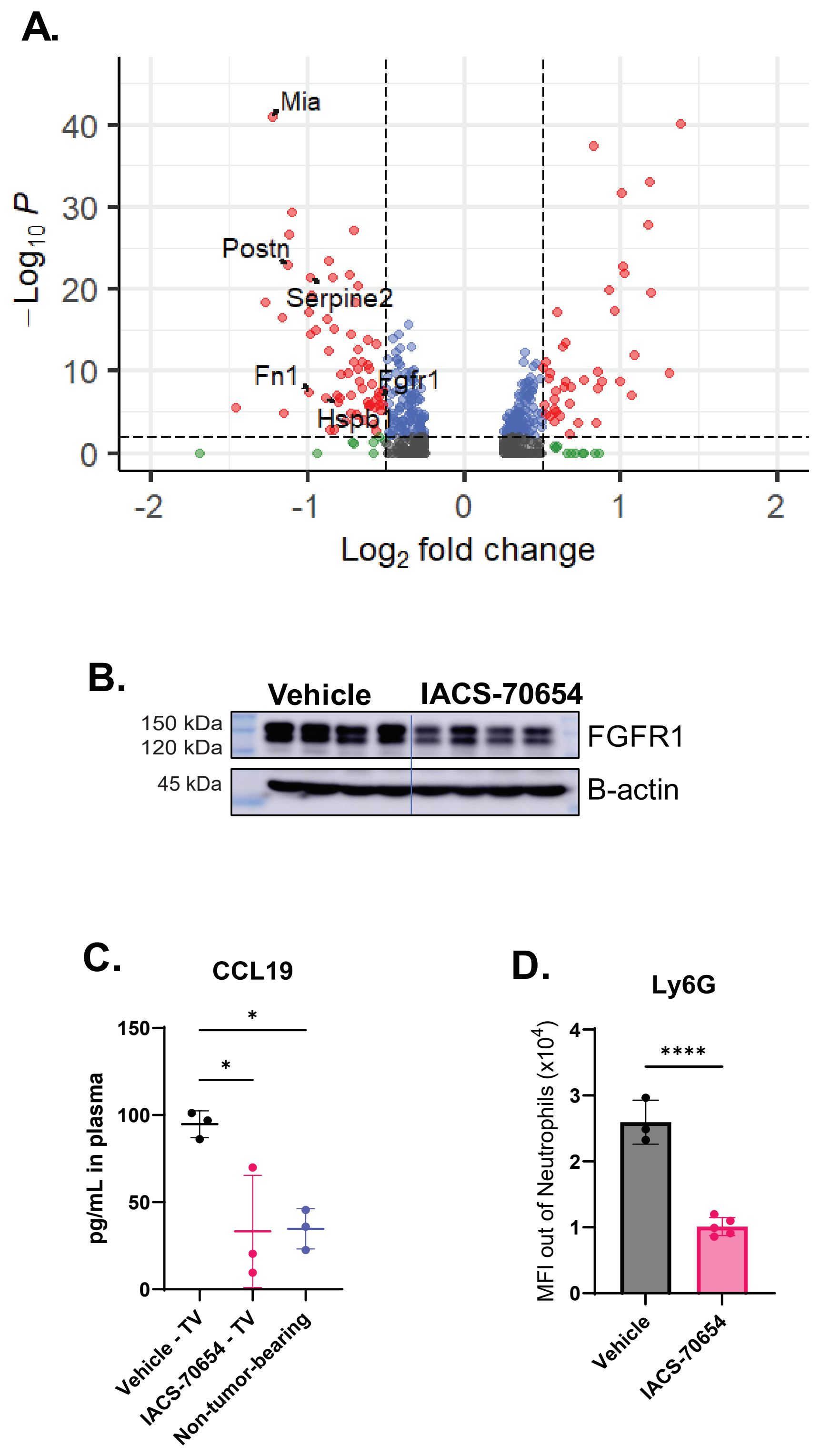

A.
Mia
Postn
Serpine2
Fn1
Fgfr1
Hspb1
$-\text{Log}_{10} P$
40
30
20
10
0
-2
-1
0
1
2
$\text{Log}_2$ fold change
B.
Vehicle
IACS-70654
150 kDa
120 kDa
45 kDa
FGFR1
B-actin
C.
CCL19
pg/mL in plasma
150
100
50
0
*
*
Vehicle - TV
IACS-70654 - TV
Non-tumor-bearing
D.
Ly6G
MFI out of Neutrophils (x10^4)
4
3
2
1
0
****
Vehicle
IACS-70654

**Supplementary Figure S1.** Structure of IACS-70654 and the weights of mice and 2208L tumors treated with IACS-70654 for 7 days. **A.** Chemical structure of IACS-70654. **B.** Mouse weights of the *Trp53*-null (2208L, T6, and T12) and PyMT-N tumor-bearing mice during the 7-day treatment of vehicle or IACS-70654. For 2208L, $n = 6$, and for all other models, $n = 5$. Error bars represent SD. **C.** Weight of 2208L, PyMT-N, T6, and T12 tumors after 7-day treatment of vehicle or IACS-70654. For all models, $n = 5$ for each treatment arm. Two-tailed unpaired Student's *t* test was used. **, $p<0.01$; ***, $p<0.001$; ****, $p<0.0001$. Error bars represent SD.

**Supplementary Figure S2.** Changes in tumor-associated myeloid cells after IACS-70654 treatment. **A.** Representative density plots of Ly6G versus Ly6C showing gating strategy for and changes in myeloid populations of T6 and T12 tumors. The gating was performed on $CD45^+/CD11b^+$ populations. **B.** Quantification of S100A8 staining on sections of 2208L tumors treated with vehicle or IACS-70654 (24, 28, or 72-hour treatment). Three representative images were analyzed for each tumor, and three biological replicates were used for each treatment arm. Two-tailed unpaired Student's *t* test was used. *, $p<0.05$. Error bars represent SD. **C.** Non-tumor-bearing WT BALB/c mice treated with vehicle or IACS-70654 for 7 days. Left: Flow cytometry analyses of blood neutrophils. Two-tailed unpaired Student's *t* test was used. Three biological replicates were used for each treatment arm. ns, $p>0.05$. Error bars represent SD. Right: Changes in mouse weight over treatment. Error bars represent SD. **D.** UMAP of TAM subpopulations with annotations in 2208L tumors treated with vehicle or IACS-70654. TAM subtypes include High GTPase expressing (GTPase), IFN response gene expressing (IFN), proangiogenic (Angio), lipid-associated (LA), resident, and proliferating (Prolif) TAMs. **E.** The fractions of TAM subtypes in TAMs of 2208L tumors treated with vehicle or IACS-70654 derived from scRNA-seq analyses. **F.** Left: Quantification of tumor-infiltrated monocytes as percentages of CD45+ cells in preclinical TNBC models treated with vehicle or IACS-70654 using flow cytometry. Monocytes are defined by $Ly6G^-/Ly6C^+$. Right: Quantification of Arginase (Arg)-positive tumor-infiltrated monocytes as percentages of $CD45^+$ cells. Two-tailed unpaired Student's *t* test was used. *, p<0.05; ***, p<0.001; ****, p<0.0001. For all models, $n \geq 5$ for each treatment arm. Error bars represent SD.

**Supplementary Figure S3.** ScRNA-seq analyses of bone marrow neutrophils. **A.** Pseudotime analysis of integrated bone marrow myeloid cells in 2208L tumor-bearing mice treated with vehicle and IACS-70654. The root is circled. **B.** Quantification of monocytes as percentages of $CD45^+$ cells in the bone marrow of 2208L tumor-bearing mice treated vehicle or IACS-70654 and non-tumor-bearing mice using flow cytometry. Monocytes are defined by $CD11b^+/Ly6G^-$. Ordinary one-way ANOVA and Tukey's multiple comparisons test were used. ****, p<0.0001; ns, p>0.05. For 2208L tumor-bearing mice, five biological replicates were used, and for the non-tumor-bearing mice, three biological replicates were used. Error bars represent SD. **C.** Quantification of G-CSF level in plasma from 2208L tumor-bearing and non-tumor-bearing mice by the cytokine/chemokine array. Three biological replicates were used for each arm. Two-tailed unpaired Student's *t* test was used. Error bars represent SD. *, p<0.05. Error bars represent SD. **D.** Volcano plot showing $-\text{Log}_{10}$ p-value versus $\text{Log}_2$ fold change in RNA expression in immature bone marrow neutrophils of 2208L tumor-bearing mice treated with IACS-70654 compared to those treated with vehicle. Genes that showed significant changes ($Log_2$ fold change > 0.5 or < -0.5 and adjusted p-value < 0.01) in expression are represented by red dots. **E.** Volcano plot showing $–Log_{10}$ p-value versus $Log_2$ fold change in RNA expression in immature bone marrow neutrophils of 2208L tumor-bearing mice treated with vehicle compared to those of non-tumor-bearing WT mice. Genes that showed significant changes ($Log_2$ fold change > 0.5 or < -0.5 and adjusted p-value < 0.01) in expression are represented by red dots. **F.** Expression distribution of *Cstdc5* and *Stfa2* in bone marrow neutrophils of 2208L tumor-bearing mice treated with vehicle or IACS-70654. **G.** GO pathway enrichment analysis of the significantly upregulated genes ($Log_2$ fold change > 0.5 and adjusted p-value < 0.01) in mature bone marrow neutrophils of 2208L tumor-bearing mice treated with IACS-70654 compared to those treated with vehicle. Biological Process (BP) gene sets from the GO database were used. The top 7 terms were listed with numbers of genes enriched.

**Supplementary Figure S4.** ScRNA-seq analyses of HSPCs. **A.** Pseudotime analysis of integrated HSPCs in 2208L tumor-bearing mice treated with vehicle and IACS-70654. The root (HSC) is circled. **B.** Volcano plot showing $–Log_{10}$ p-value versus $Log_2$ fold change in RNA expression in cluster 4 of HSPCs in the bone marrow of 2208L tumor-bearing mice treated with IACS-70654 compared to those treated with vehicle. Genes that showed significant changes ($Log_2$ fold change > 0.5 or < -0.5 and adjusted p-value < 0.01) in expression are represented by red dots. **C.** Violin plots showing the RNA expression of *Txnip* and *Ifitm3* in HSCs of 2208L tumor-bearing mice treated with vehicle or IACS-70654 and non-tumor-bearing WT mice. **D.** Violin plots showing the RNA expression of *Txnip* and *Ifitm3* in MPPs of 2208L tumor-bearing mice treated with vehicle or IACS-70654 and non-tumor-bearing WT mice. **E.** Violin plots showing the

RNA expression of *Calr* in CMP-1 and MPPs of 2208L tumor-bearing mice treated with vehicle or IACS-70654 and non-tumor-bearing WT mice. **F.** Expression distribution of *Ifitm1* in bone marrow HSPCs of 2208L tumor-bearing mice treated with vehicle or IACS-70654 and non-tumor-bearing WT mice. **G.** Quantification of IL-3 level in plasma collected from 2208L tumor-bearing mice treated with vehicle or IACS-70654 by the cytokine/chemokine array. Three biological replicates were used for each treatment arm. Two-tailed unpaired Student's *t* test was used. Error bars represent SD. *, $p<0.05$. Error bars represent SD.

**Supplementary Figure S5.** ScRNA-seq analyses of 2208L tumors treated with vehicle or IACS-70654. **A.** UMAP of integrated 2208L tumor samples treated with vehicle and IACS-70654 with cell type annotations. **B.** Expression distribution of *Krt18, Krt8, Epcam,* and *Ptprc* in integrated 2208L tumor samples. **C.** GO pathway enrichment analysis of the significantly downregulated genes ($Log_2$ fold change < -0.5 and adjusted p-value < 0.01) in 2208L tumor cells treated with IACS-70654 compared to those treated with vehicle. Biological Process (BP) gene sets from the GO database were used. The top 10 terms were listed with numbers of genes enriched.

**Supplementary Figure S6.** Combination treatment of IACS-70654 with DTX and anti-PD-1 and anti-CD8. **A.** Quantification of CTLs as percentages of CD45+ cells in 2208L tumors treated vehicle or IACS-70654 with or without CTL depletion using flow cytometry. Ordinary one-way ANOVA and Tukey's multiple comparisons test were used. For all groups, more than four biological replicates were used. ****, $p<0.0001$; *, $p<0.05$. Error bars represent SD. **B.** Flow cytometry analysis of infiltrated PD-1+ CTL in the 2208L tumors treated with vehicle or IACS-70654 for 18 days. For each group, five biological replicates were used. Two-tailed unpaired Student's *t* test was used. ns, p>0,05. Error bars represent SD. **C.** Changes in mouse weight of 2208L tumor-bearing mice of all treatment groups over days of treatment. Error bars represent SD. **D.** Tumor growth curves of 2208L tumors of all treatment groups.

**Supplementary Figure S7.** Analysis of metastasis-related genes in 2208L tumors and 2208L lung metastases. **A.** Volcano plot showing –$Log_{10}$ p-value versus $Log_2$ fold change in RNA expression in the tumor cells of 2208L tumors treated with IACS-70654 compared to those treated with vehicle. Genes that showed significant changes ($Log_2$ fold change > 0.5 or < -0.05 and adjusted p-value < 0.01) in expression are represented by red dots. The genes that are associated with tumor migration, invasion, and metastasis are labeled. **B.** Immunoblotting analysis of FGFR1 expression in 2208L tumors treated with vehicle or IACS-70654 for 7 days. For each treatment arm, four biological replicates were used. β-actin is used as the loading control. **C.** Quantification of CCL19 level in plasma from 2208L lung metastases-bearing mice treated with vehicle or IACS-70654 and non-tumor-bearing mice by the cytokine/chemokine array. Three biological replicates were used for each treatment arm. Ordinary one-way ANOVA and Tukey's multiple comparisons test were used. *, p<0.05. Error bars represent SD. **D.** Flow cytometry analyses showing median fluorescent intensity (MFI) of Ly6G in blood neutrophils of 2208L lung metastases-bearing mice treated with vehicle or IACS-70654. Two-tailed unpaired Student's *t* test was used. Error bars represent SD. ****, p<0.0001. Error bars represent SD.
